# Mechanistic dissection of a dopamine-gated cation channel from *Daphnia* reveals key determinants of ligand selectivity and sensitivity

**DOI:** 10.1101/2025.10.31.685654

**Authors:** Tina M. McRunnel, Thomas E. Reynoldson, Yu Zhu, Taufiq Rahman, Iris Hardege

## Abstract

The pentameric ligand-gated ion channels (pLGICs) are a deeply conserved superfamily that includes nicotinic acetylcholine (nAChR), GABA_A_Rs, glycine, and serotonin receptors and mediates fast neurotransmission. While vertebrate pLGICs have been extensively characterized, recent discoveries in invertebrates have revealed atypical members with novel ligand specificities, including dopamine-gated ion channels (dop-LGICs) that enable direct ionotropic dopaminergic signaling. Here, we investigate the molecular determinants of catecholamine sensitivity and selectivity in a cation-permeable dopamine-gated receptor from *Daphnia magna* (Dm-DopC1), which shares high structural homology with vertebrate nAChRs. We find that mutations in loop B (D188N) near-abolish catecholamine responses, while substitutions in loops A (S123P) and C (S227G, E230P) significantly alter sensitivity and ligand selectivity, highlighting the cooperative role of these loops in shaping receptor pharmacology. Complementary face mutations in loops D and E (N92T, A158G) further refined dopamine versus norepinephrine sensitivity. Finally, we also show that this dopamine receptor can be antagonized by both nicotinic and dopaminergic compounds, underscoring its hybrid pharmacological profile. Together, these results reveal how subtle changes in conserved binding motifs can shift receptor specificity from cholinergic to dopaminergic signaling, providing new insights into the evolutionary flexibility of pLGICs and the molecular basis of ionotropic catecholamine transmission in invertebrates.

## 1. Introduction

The processes governing neural communication are dynamic and highly evolutionarily conserved in animals across phyla (Moroz et al., 2021). Relying on the highly orchestrated molecular mechanisms that mediate the release and detection of neurotransmitters, converting extracellular chemical cues into either rapid electrical shifts in cell membrane potential or slower intracellular signaling cascades. Neurotransmission occurs via two distinct pathways: ionotropic and metabotropic. Ionotropic receptors, mediate fast synaptic transmission while in contrast, metabotropic receptors act through slow and second messenger pathways and produce longer lasting, modulatory effects.

The most widespread ionotropic receptors in animals fall into the pentameric or Cys-loop ligand gated ion channel superfamily (pLGIC), which includes nicotinic acetylcholine receptors (nAChR), GABA_A_ receptors, glycine receptors (GlyR) and serotonin receptors (5HT_3_R), each typically associated with distinct neurotransmitters and ionic conductances. The structural architecture of pLGICs is highly conserved and consists of five subunits arranged symmetrically around a central ion pore (Figure 1A). Each subunit contributes a β-sandwich extracellular domain (ECD) and four transmembrane α-helices. Neurotransmitter binding occurs at the interfaces between subunits, where one subunit serves as the ‘principal face’ contributing three extracellular loops A-C and the other contributes loops D-F to form a composite ligand binding site (Figure 1B) (Lynagh and Pless, 2014). While some pLGICs function as homomers most receptors are formed of heteromeric combinations of up to 5 different subunits (Boulin et al., 2008) with the ligand binding interface only forming between specific subunit combinations. Despite this conserved scaffold, structural studies have revealed how small differences in loop composition and conformation underlie substantial diversity in ligand recognition and receptor pharmacology (Corringer et al., 2010; Dalal et al., 2025; Dani, 2015; Pan et al., 2012; Sauguet et al., 2014; Unwin, 2003). This class of receptors have been extensively studied in vertebrate systems, and these receptors have become major pharmacological targets, forming the basis of anesthetics, antiparasitic and anxiolytics. This vertebrate-centric knowledge base has shaped much of our understanding of synaptic transmission. However, studies in invertebrate models have uncovered atypical pLGICs with unexpected ligand specificity, gating properties and synaptic roles (Cully et al., 1996; Ringstad et al., 2009). Characterizing these divergent receptors is essential for understanding the evolutionary diversity and functional plasticity possible in neurotransmission. For example, it was once assumed that each neurotransmitter exerts either an excitatory or an inhibitory effect depending on the receptor class that it activates. However, the discovery of both excitatory and inhibitory glutamate-gated chloride channels (GluCls) in *Caenorhabditis elegans* and *Drosophila melanogaster* challenged this view (Cully et al., 1996, 1994). These invertebrate-specific channels are the targets for the antiparasitic treatment ivermectin, which locks the channel in an open state, causing paralysis in parasitic worms.

**Figure 1.**
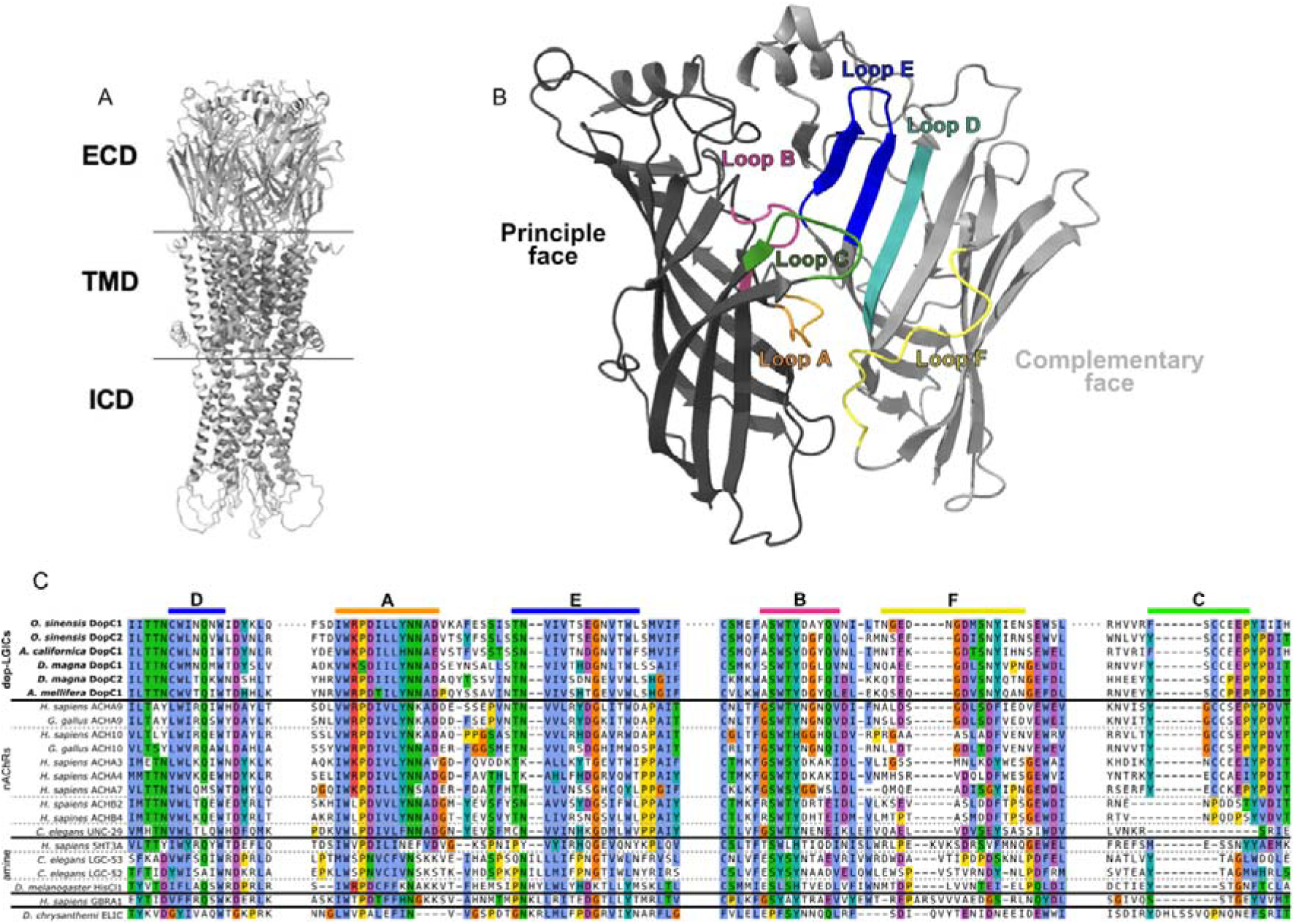
**A.** Full length AlphaFold2 model of Dm-DopC1 with the extracellular (ECD), transmembrane (TM) and intracellular (ICD) domains highlighted. **B.** Close up of the ECD and ligand binding domain made up of the interface between two adjacent subunits (light and dark grey). Extracellular loops associated with ligand binding are labeled as follows; loop A (orange), loop B (pink), loop C (green), loop D (turquoise), loop E (blue), loop F (yellow). **C.** Trimmed alignment of the ECD of dop-LGICs, nAChRs and other pLGICs showing the ligand binding loops. Alignment generated with MAFFT and residues colored with CLUSTAL coloring.

One of the most intriguing discoveries in invertebrate systems is the identification of ligand-gated ion channels gated by dopamine (dop-LGICs), which enable direct, ionotropic dopaminergic signalling (Courtney et al., 2025; Mitchell et al., 2025; Morud et al., 2021; Ringstad et al., 2009). This mode of neurotransmission contrasts with canonical dopaminergic signalling in mammals, which is mediated by GPCRs. The most recently identified cation permeable dop-LGIC which is gated by the catecholamines dopamine, norepinephrine and epinephrine has been shown to be conserved in multiple invertebrate species, including *Daphnia magna, Apis mellifera*, *Aplysia california* and *Octopus sinensis* (Courtney et al., 2025; Mitchell et al., 2025). In contrast, the anion dop-LGICs studied previously appear to be nematode specific and gated by dopamine and to a lesser extent tyramine (Morud et al., 2021; Ringstad et al., 2009). Intriguingly the invertebrate cation dop-LGIC shares close sequence homology with the vertebrate α9α10 nicotinic acetylcholine receptor (nAChR), including within the classic ligand binding domains. A great deal of previous research, including more recent structural studies have revealed residues that contribute to the binding of cholinergic ligands for nAChRs (Dani, 2015; Morales-Perez et al., 2016; Su et al., 2025); however, we found that many of these are conserved in cationic dop-LGICs (Figure 1C). In this study we sought to unravel the contribution of putative ligand binding domains and individual residues in the homomeric *Daphnia magna* dop-LGIC, Dm-DopC1, in the sensitivity of cationic dop-LGICs to catecholamines rather than cholinergic ligands like its close relatives the nAChRs.

## 2. Materials and methods

### 2.1 Structural Modelling and Sequence Alignments

Amino acid sequences of a range of pLGICs (full sequences are provided in Table S 1) were aligned using the MAFTT algorithm and visualized with SnapGene (Dotmatics). Structural models were predicted using AlphaFold ColabFold. For *D. magna* Dm-DopC1, five predicted models were generated and overlaid in PyMOL. Residue positions were evaluated for consistency and model confidence (pLDDT > 70). Sites exhibiting conserved structural placement and high-confidence predictions were considered for site-directed mutagenesis.

### 2.2 Molecular Biology and Mutagenesis

Mutations were introduced using the Q5® Site-Directed Mutagenesis Kit (New England Biolabs). Constructs were linearized using NotI. cRNA was synthesized using the T3 mMessage mMachine Kit (ThermoFisher) and purified with the GeneJET RNA Cleanup Kit (ThermoFisher Scientific).

### 2.3 *Xenopus laevis* Oocyte Preparation and RNA Injection

Stage V-VI defolliculated *Xenopus laevis* oocytes were obtained from EcoCyte Bioscience and incubated in ND96 solution (in mM): 96 NaCl, 2 KCl, 1.8 CaCl2, 1 MgCl2, 5 HEPES, pH 7.5. Oocytes were stored at 18°C and injected within two days of delivery. For RNA injection, oocytes were placed in 96-well plates and injected using the Roboinject system (Multi Channel Systems, Germany). Each oocyte was injected with 50 nL of cRNA (250ng/µL). Oocytes were incubated at 18°C for 1 day prior to electrophysiological recordings.

### 2.4 Two-Electrode Voltage Clamp (TEVC) Recordings

TEVC recordings were performed using the Robocyte2 automated platform (Multi Channel Systems, Germany). Microelectrodes (0.5-1.5 MΩ resistance) were pulled using a P-97 micropipette puller (Sutter Instrument) and filled with 1 M KCl and 1.5 M acetic acid. Oocytes were voltage-clamped at -60 mV and recordings were sampled at 500 Hz throughout. Agonist solutions (dopamine, norepinephrine, epinephrine, tyramine, acetylcholine and serotonin) were prepared in ND96 on the day of recording and applied at concentrations ranging from 0.1 µM to 10 mM. Each ligand was perfused for 10 s followed by a 60 s wash. Wild-type oocytes were included in every recording batch. For antagonist experiment, compound solutions (50 nM to 500 μM, unless otherwise stated) were prepared in ND96 on the day of recording with 50 µM dopamine (Dm-DopC1). Before and after compound application, response to dopamine-only was recorded to obtain an Imax value per oocyte and to screen for potential irreversible binding of the antagonist. All compounds were perfused for 7 s followed by a 60 s wash. Perfusion speed was set to 3 ml/min throughout. Compounds used as antagonists: Acetylcholine chloride (Sigma-Aldrich (A6625)), Apomorphine hydrochloride (Abcam (ab269887)), Atropine (Abcam (ab145582)), Isoprenaline hydrochloride (Sigma-Aldrich (I5627)), Levamisole hydrochloride (MedChemExpress (HY-13666)), (-)-Nicotine detartrate (Abcam (ab120562)), Propranolol hydrochloride (Santa-Cruz Biotechnology (sc-3580)), (+)-Tubocurarine chloride pentahydrate (Sigma-Aldrich (93750)).

### 2.5 Data Analysis

All current amplitudes were exported using the Roboocyte2+ software (Multi Channel Systems) and normalized per oocyte to the maximum response (I/Imax) to account for variation in expression levels using a custom python script adapted from (Morud et al., 2021). Outliers were identified for each construct and removed within GraphPad Prism 10 using the ROUT method (Q=1%) for every individual agonist/antagonist concentration. Agonist and antagonist dose response curves were fitted in GraphPad Prism 10 using a three-parameter logistic equation with a fixed Hill Slope of 1, unless otherwise stated. EC_50_ and IC_50_ values were calculated from the fitted curves and reported as mean *±* standard error of mean (SEM). Statistical significance between wild-type and mutant dose-response curves was assessed using a sum-of-squares F test. Peak current values were extracted using an existing Python script (Morud et al., 2021). Normalized peak currents from the panel of ligands (1mM each: tyramine, acetylcholine, norepinephrine, serotonin and epinephrine) were analyzed using a one-way ANOVA followed by a Fisher’s Least Significant Difference (LSD) post hoc test. All TEVC recordings were baseline-subtracted unless otherwise stated. The same pooled wild-type data was used for all comparisons.

### 2.6 *In silico* prediction of Dm-DopC1 protein in complex with catecholamines and acetylcholine

To predict the possible binding mode of different ligands onto Dm-DopC1 protein (GenBank accession no. XP_032790382.1), we used AlphaFold3-based co-folding approach (Abramson et al., 2024). The N-terminal signal peptide sequence (residues 1-26) was excluded prior to modelling, and the remaining sequence was used to model as homodimer whilst each of the small-molecule ligands (dopamine, norepinephrine, epinephrine and acetylcholine) was incorporated directly into the modelling process using their corresponding Chemical Component Dictionary (CCD) codes. For each ligand, the protein dimer and a single ligand molecule were provided simultaneously as inputs, enabling joint prediction of protein conformation and ligand binding pose within a single inference step. Five independent structures were generated for each protein–ligand combination using an identical protocol to ensure methodological consistency and to account for stochastic variation in model sampling. The best model was considered for each ligand and the corresponding 2D ligand interaction diagram was generated using PoseView^™^ (Stierand and Rarey, 2010) accessed through the Protein Plus server (https://proteins.plus/).

### 2.7 Software and Code Availability

TEVC data were analyzed using custom Python scripts (https://github.com/hiris25/TEVC-analysis-scripts). GraphPad Prism 10 was used for data plotting and fitting. MAFFT (Katoh and Standley, 2013) was used for sequence alignment, alignments were visualized in snapgene and ChimeraX 1.9 and Pymol were used for 3D structural visualization. All constructs and raw data are available upon request.

## 3. Results

### 3.1 Despite strong conservation with nAChRs, residues in loop C contribute to catecholamine sensitivity and selectivity

The principle or α-type subunit contributes three structurally distinct loops to the ligand binding domain, with loop C forming the front outer side of the binding pocket that closes around the ligand during receptor activation and loops A and B forming the back of the binding pocket (Dani, 2015; Morales-Perez et al., 2016). Loop C contains two conserved structural features in nAChRs. The first is a pair of tyrosines implicated in aromatic cage formation (Braun et al., 2016; Celie et al., 2004; Tomaselli et al., 1991) and the second is a double cystine motif (Tomizawa et al., 2008) together forming a ‘YXCC’ motif. Both features are present in dop-LGICs and in Dm-DopC1 are represented by Y226 and Y233, and the C228 - C229, respectively (Figure 2E-F). Notably though both human and chicken nAChR α9 and α10 subunits contain a glycine at the ‘X’ position, while the dop-LGICs contain serine at this position. When we substituted S227 for glycine at this position in Dm-DopC1 we observed a significant reduction in sensitivity to all catecholamines (Figure 2A-C). The largest shift was seen with dopamine, shifting 7-fold from a WT EC_50_ of 84µM to 589µM, norepinephrine EC_50_ also shifted 4-fold and epinephrine 1.8-fold (Figure 2A-C). This suggests that the native serine may stabilise local interactions required for maintaining the C loop conformation or supporting neighbouring aromatic residues for catecholamine recognition.

**Figure 2.**
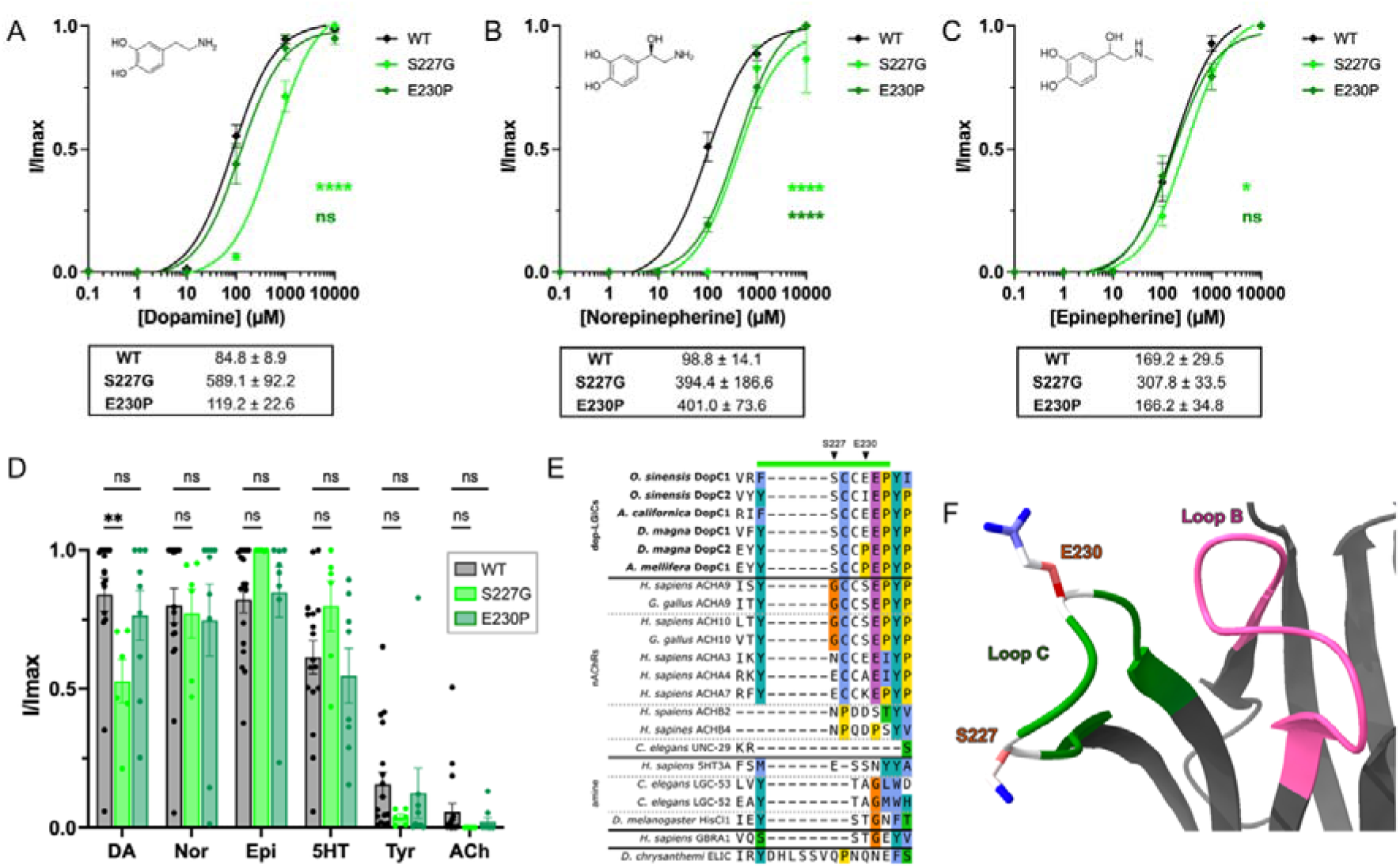
Mutations in Loop C of a *D. magna* dop-LGIC reduce catecholamine sensitivity and alter agonist selectivity. **A-C.** Dose response curves for dopamine (A), norepinephrine (B) and epinephrine (C) in wild-type, S227G and E230P Dm-DopC1 expressing *Xenopus* oocytes. Current normalized by I/Imax. EC_50_ values (± SEM) shown below each plot. **** = P<0.0001, * = P<0.05, ns = not significant calculated by sum of squares F test. WT: *n* = 24 (dopamine), 12 (norepinephrine), 11 (epinephrine); S227G: *n* = 6 (dopamine), 3 (norepinephrine), 5 (epinephrine); E230P: *n* = 10 (dopamine), 4 (norepinephrine), 9 (epinephrine). **D.** Relative maximum I/Imax currents of *Xenopus* oocytes expressing wild-type, S227G and E230P Dm-DopC1, perfused for 10 s with 1 mM each: dopamine (DA), norepinephrine (Nor), epinephrine (Epi), serotonin (5HT), tyramine (Tyr), and acetylcholine (ACh). WT: *n* = 18 (dopamine, norepinephrine, ACh, tyramine), 17 (epinephrine, serotonin); S227G: *n* = 11 per agonist; E230P: *n* = 6 per agonist. ** = P<0.01, ns = not significant by one-way ANOVA and a Fisher’s LSD. **E.** Sequence alignment of the Loop C region. Residues S227 and E230 indicated by arrows, loop C is highlighted by the green bar. **F.** AlphaFold2 model of the loop C (green) region of Dm-DopC1 with S227 and E230 relative to Loop B (pink) at the ligand-binding site.

We were also interested in the residue immediately following the ‘YXCC’ motif, which is divergent across species and receptor types. However, both Dm-DopC2 and Am-DopC1 (from *Apis mellifera*), which have a higher relative selectivity for dopamine vs. the other catecholamines, encode a proline at this position. A substitution to proline in this position in Dm-DopC1 (E230P) produced a selective decrease in norepinephrine sensitivity only, with a 4-fold shift of EC_50_ from 98µM to 401µM, while maintaining wild type sensitivity for dopamine and epinephrine (Figure 2A-C). However, despite altering the relative sensitivities, the E230P mutation did not alter the ligand binding profile of the receptor, with norepinephrine still eliciting similar relative peak currents to dopamine and epinephrine at 1mM (Figure 2D). In contrast S227G which altered sensitivity to all catecholamines, abelite with more pronounced shift in dopamine sensitivity, did alter the ligand binding profile such that dopamine induced currents at 1mM fell significantly to approximately 50% of the maximum currents achieved with norepinephrine and epinephrine (Figure 2D). Together these data suggest that despite key nAChR features of loop C being conserved between cholinergic and catecholamine binding receptors, the surrounding residues also contribute to ligand specificity and binding capability.

### 3.2 A conserved aspartate in loop B of principle face is critical to catecholamine sensitivity

Loop B forms part of the back wall of the ligand binding domains on the principle face and contains one of the conserved tryptophans forming the aromatic cage which is part of a conserved region encoded the consensus motif ‘SWTY’ (Figure 3E-F). A semi conserved glycine, 4 amino acids downstream of the tryptophan has been implicated in defining high and low potency nAChRs (Puskar et al., 2012). In the dop-LGICs this post-tryptophan glycine is broadly conserved and directly preceded by an aspartate which is conserved across the dop-LGICs but not found in vertebrate α9/10 receptors. Substitution of this position (D188) in Dm-DopC1 with asparagine, more commonly found in nAChRs, led to a striking reduction in catecholamine sensitivity, leading to EC_50_ values for all three ligands of over 9mM, although true EC_50_ values could not be calculated due to lack of saturation even at 10mM (Figure 3A-C). The D188N mutation also led to significant changes in ligand binding profiles at 1mM with norepinephrine currents significantly reduced to 8% of the maximal currents, dopamine reduced to 37% and serotonin induced currents also falling to 19% (Figure 3D). However, it is worth noting that there was no appreciable difference in acetylcholine currents, which along with the residual currents observed from the catecholamines suggests that while loop B is critical to catecholamine gating, this single residue switch is insufficient in defining catecholamine vs cholinergic activation.

**Figure 3.**
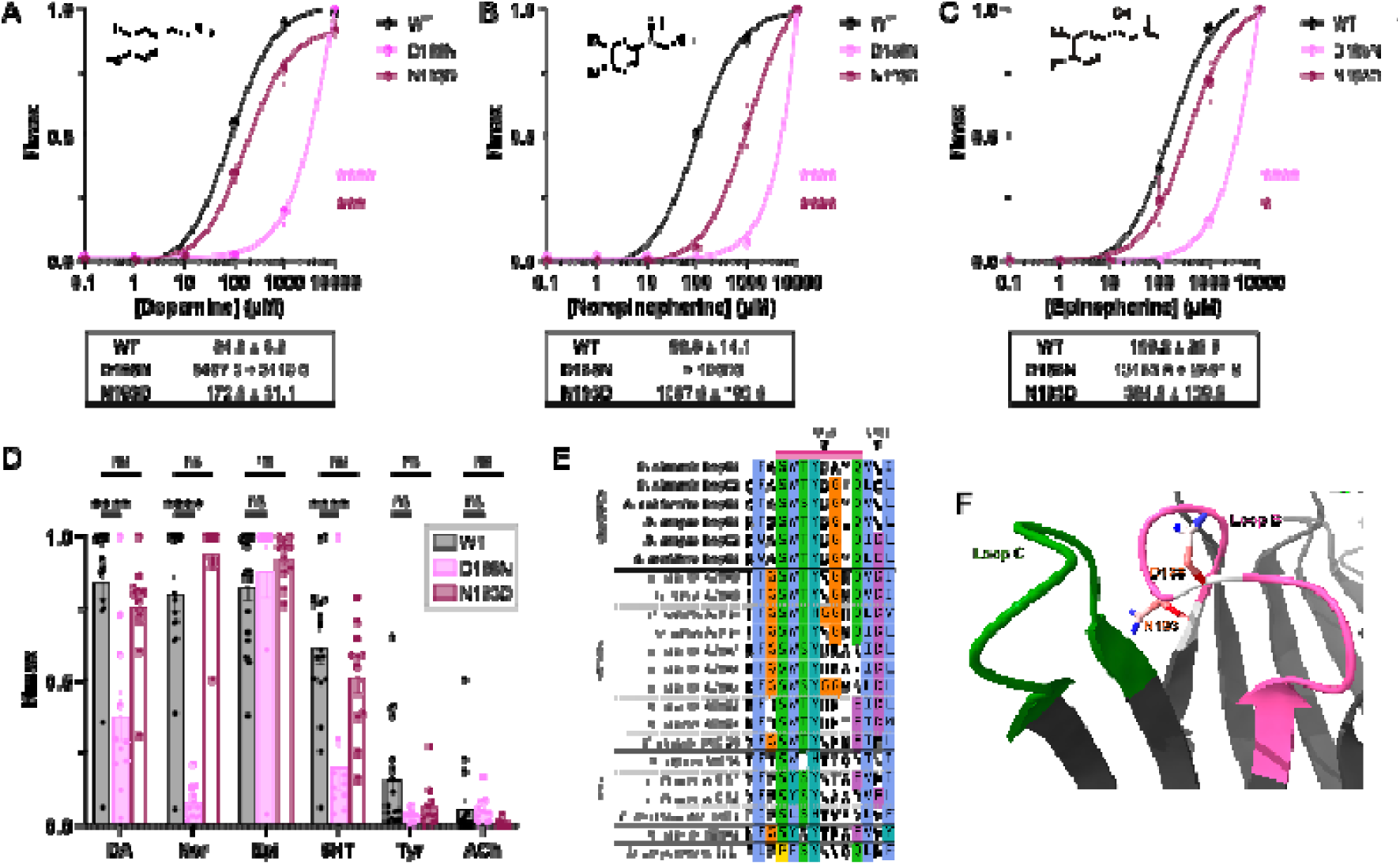
Mutations in Loop B of Dm-DopC1 disrupt catecholamine sensitivity and alter ligand selectivity. **A-C.** Dose response curves for dopamine (A), norepinephrine (B) and epinephrine (C) in wild-type, D188N and N193D Dm-DopC1 expressing *Xenopus* oocytes. Current normalized by I/Imax. EC_50_ values (± SEM) shown below each plot. **** = P<0.0001, *** = P<0.001, * = P<0.05 calculated by sum of squares F test. WT: *n* = 24 (dopamine), 12 (norepinephrine), 11 (epinephrine); D188N: *n* = 10 (dopamine), 5 (norepinephrine), 9 (epinephrine); N193D: *n* = 15 (dopamine), 5 (norepinephrine), 7 (epinephrine). **D.** Relative maximum I/Imax currents of *Xenopus* oocytes expressing wild-type, D188N and N193D Dm-DopC1, perfused for 10 s with 1 mM each: dopamine (DA), norepinephrine (Nor), epinephrine (Epi), serotonin (5HT), tyramine (Tyr), and acetylcholine (ACh). WT: *n* = 18 (dopamine, norepinephrine, ACh, tyramine), 17 (epinephrine, serotonin); D188N: *n* = 11 per agonist; N193D: *n* = 11 per agonist. **** = P<0.0001, ns = not significant by one-way ANOVA and a Fisher’s LSD. **E.** Sequence alignment of the Loop B region. Residues D188 and N193 indicated by arrows, loop B is highlighted by the pink bar. **F.** AlphaFold2 model of the loop B (pink) region of Dm-DopC1 with D188N and N193D relative to Loop C (green) at the ligand-binding site.

We also noted an additional residue of interest downstream of loop B, which differed only in Dm-DopC1 compared to other dop-LGICs and vertebrate α9/10. Given the differences in selectivity between Dm-DopC1 and Dm-DopC2 we hypothesised that the conserved aspartate at this position may also be contributing to ligand selectivity. Mutation at this location in Dm-DopC1 (N193D), led to a significant increase in EC_50_ for all catecholamines, most notably for norepinephrine with a 11-fold shift resulting in an EC_50_ of 1087μM, with smaller yet statistically significant shifts for dopamine and epinephrine of approximately 2-fold (Figure 3A-C). There was no change in overall ligand binding profile with all 3 catecholamines still eliciting the largest currents (Figure 3D). This suggests that the conserved aspartate at this position may contribute to selectivity between catecholamines, with receptors bearing this residue being relatively more selective for dopamine.

### 3.3 Residues on the complementary face of the binding pocket contribute to dopamine sensitivity and selectivity

In addition to the principal subunit, two loops on the complementary face of the binding pocket have also been shown to be involved in ligand binding in nAChRs. Loop D is composed of a single β strand and forms the base of the binding site, while loop E is composed of two β strands connected by a short loop. We observed that loop D was broadly conserved between both nAChRs and dop-LGICs, however we again noted that the two dop-LGICs with higher dopamine selectivity, Dm-DopC2 and Am-DopC1, encoded a threonine in the centre of the loop (Figure 4E-F). Creating this change in Dm-DopC1, N92T, led to a striking and significant reduction of norepinephrine sensitivity, with an EC_50_ increase from 98μM to 439μM, with no effect on dopamine or epinephrine (Figure 4A-C) and no changes to overall ligand binding profile at 1mM (Figure 4D).This suggests that the threonine at this position is important for discerning between norepinephrine and the other catecholamines.

**Figure 4.**
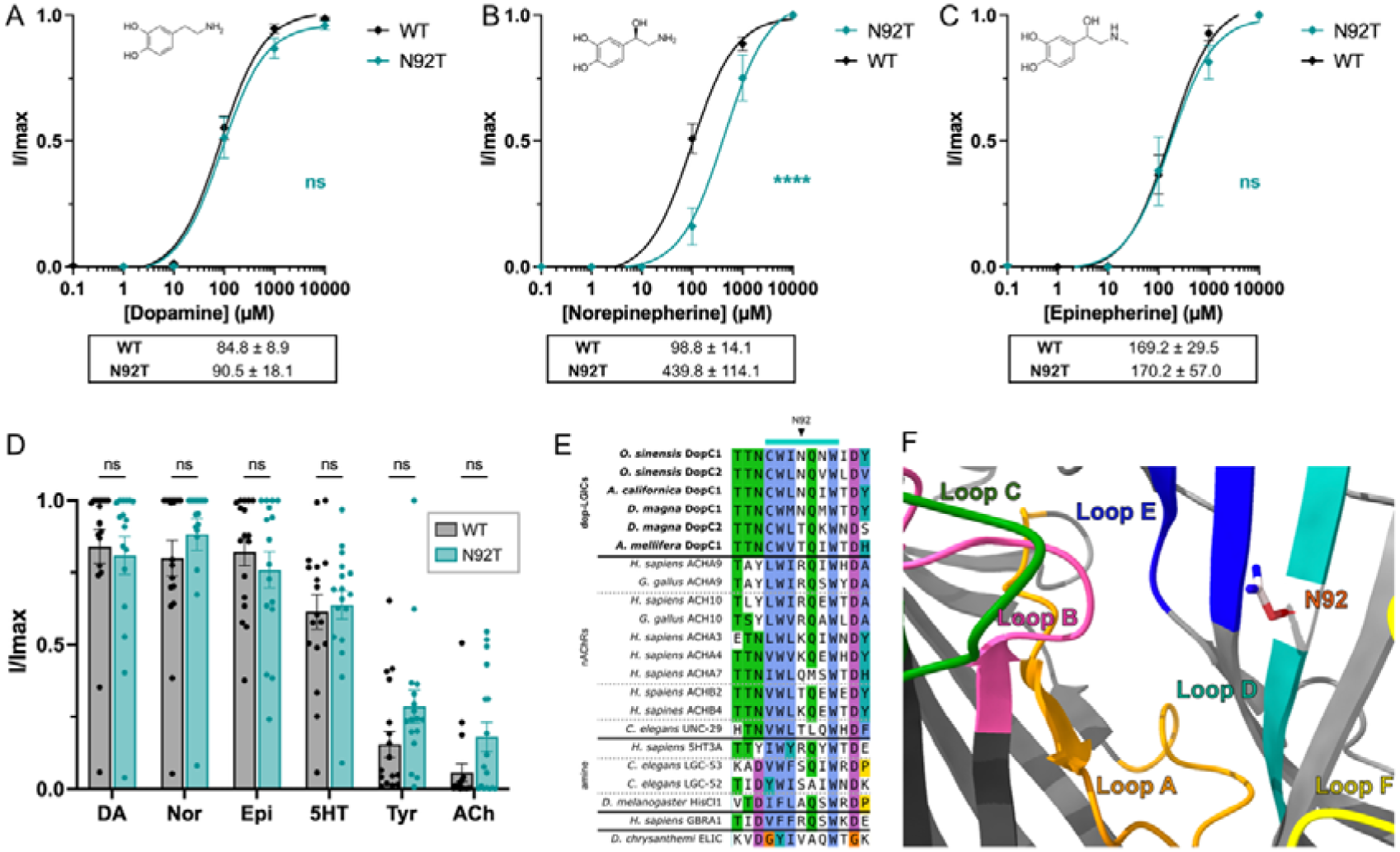
The N92T mutation in Loop D of a *D. magna* dop-LGIC reduces norepinephrine sensitivity but preserves agonist selectivity. **A-C.** Dose response curves for dopamine (A), norepinephrine (B) and epinephrine (C) in wild-type and N92T Dm-DopC1 expressing *Xenopus* oocytes. Current normalized by I/Imax. EC_50_ values (± SEM) shown below each plot. **** = P<0.0001, ns = not significant calculated by sum of squares F test. WT: *n* = 24 (dopamine), 12 (norepinephrine), 11 (epinephrine); N92T: *n* = 10 (dopamine), 5 (norepinephrine), 8 (epinephrine). **D.** Relative maximum I/Imax currents of *Xenopus* oocytes expressing wild-type and N92T Dm-DopC1, perfused for 10 s with 1 mM each: dopamine (DA), norepinephrine (Nor), epinephrine (Epi), serotonin (5HT), tyramine (Tyr), and acetylcholine (ACh). WT: *n* = 18 (dopamine, norepinephrine, ACh, tyramine), 17 (epinephrine, serotonin); N92T: *n* = 11 per agonist. ns = not significant by one-way ANOVA and a Fisher’s LSD. **E.** Sequence alignment of the loop D region. Residue N92 is indicated by an arrow, loop D is highlighted by the turquoise bar. **F.** AlphaFold2 model of the loop D (turquoise) region of Dm-DopC1 with N92 relative to loop B (pink), loop C (green), loop F (yellow), loop A (orange) and loop E (blue) at the ligand-binding site.

Despite overall high similarity within loop E, sequence alignments revealed a strongly conserved aspartate at the proximal end of loop E in α9/10 that is missing from the dop-LGICs, which frequently encode a hydrophobic leucine or phenylalanine at this position (Figure 4E-F). When we substituted leucine with aspartate (L155D) at this location in Dm-DopC1 we could not generate functional receptors that responded consistently to any of the ligands tested, although due to leak current in oocytes we hypothesise that the protein was expressed (Figure S 1G-H). This suggests that leucine at this position is critical to either channel formation or ligand binding. Additionally, we also noted that the two more-dopamine selective dop-LGICs, Dm-DopC2 and Am-DopC1, had a substitution of an alanine conserved across both receptor types with a glycine (Figure S 1E-F). Despite being a minor alteration in sequence we found that this substitution in Dm-DopC1, A158G, led to small but significant increases in EC_50_ for all catecholamines to similar extents (EC_50_s: dopamine: 235μM, norepinephrine: 440μM, epinephrine: 438μM) (Figure S 1A-D). Again, supporting the notion that residues surrounding key ligand binding loops contribute to receptor function, likely through the stabilisation of these regions.

### 3.4 The conserved proline of loop A enhances catecholamine sensitivity

In line with previous findings, our sequence alignments identified a proline residue conserved across all pLGICs that sits within the conserved ‘WXPD’ motif (Figure 5E-F) (Braun et al., 2016). However, in Dm-DopC1 this location encodes a serine, which AlphaFold2 structural modelling placed near the inter-subunit ligand-binding interface, in proximity to Loops B and C (Figure 5F). Strikingly, restoring the proline at this position in Dm-DopC1 (S123P) significantly increased sensitivity to all catecholamines (Figure 5A-C). Dopamine EC_50_ decreased from 84.82D±D9.1DµM to 39.84D±D6.06DµM, norepinephrine from 98.78D±D14.09DµM to 36.52D±D7.86DµM, and epinephrine from 169.17D±D29.5DµM to 47.87D±D14.05DµM (Figure 5A-C). Interestingly, unlike the other mutations examined in this study, S123P also led to an increase in Hill slope for norepinephrine (2.16D±D0.19) compared to WT (1.31D±D0.30), suggesting enhanced co-operativity (Figure S 2). Overall, the relative ligand binding profile was unaltered, although there was a small but non-significant increase in relative acetylcholine current (Figure 5D).

**Figure 5.**
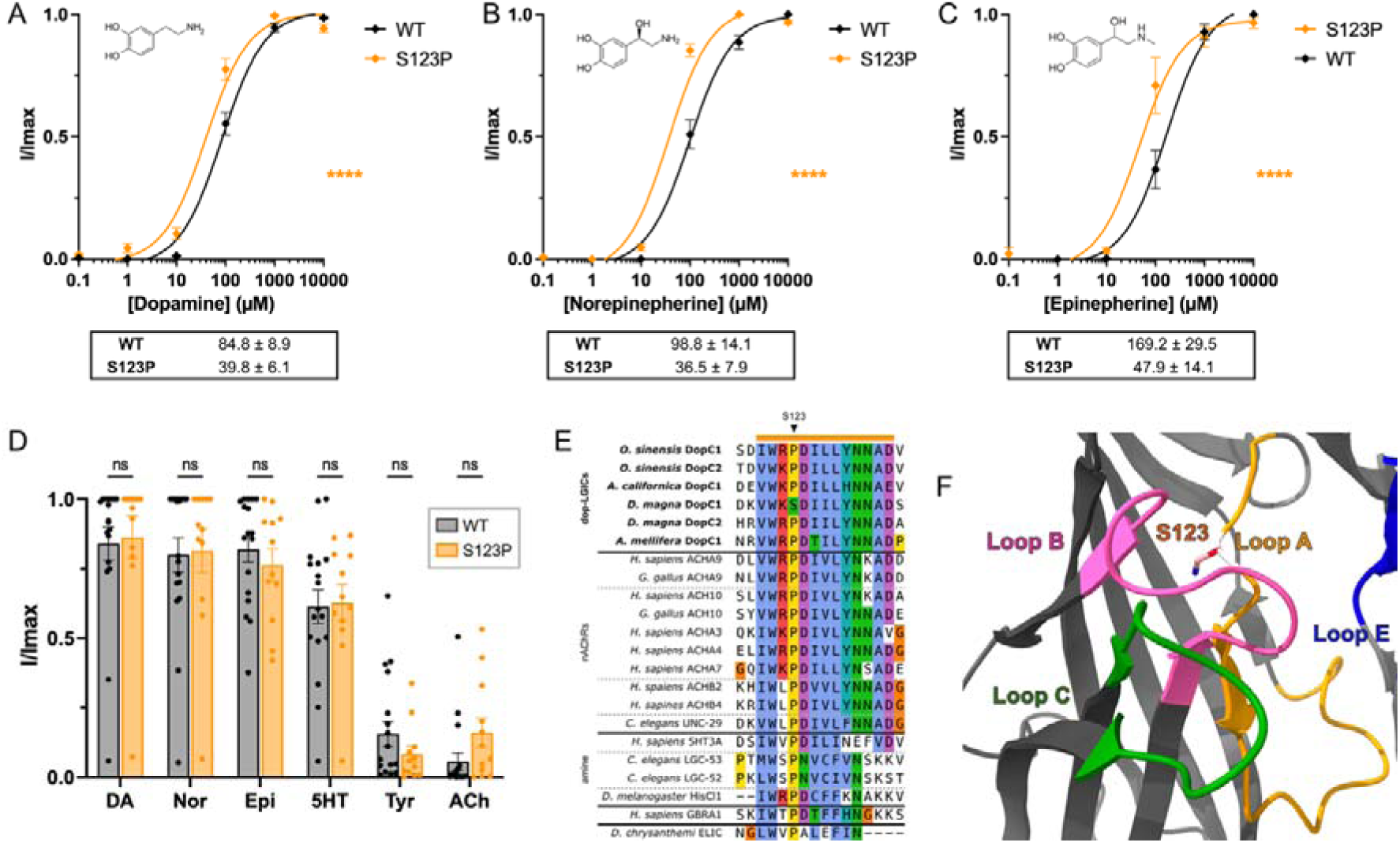
The conserved proline of loop A is required for ligand sensitivity in dop-LGICs. **A-C.** Dose response curves for dopamine (A), norepinephrine (B) and epinephrine (C) in wild-type and S123P Dm-DopC1 expressing *Xenopus* oocytes. Current normalized by I/Imax. EC_50_ values (± SEM) shown below each plot. **** = P<0.0001 calculated by sum of squares F test. WT: *n* = 24 (dopamine), 12 (norepinephrine), 11 (epinephrine); S123P: n = 12, (dopamine), 4 (norepinephrine) and 8 (epinephrine). **D.** Relative maximum I/Imax currents of *Xenopus* oocytes expressing wild-type and A158G Dm-DopC1, perfused for 10 s with 1 mM each: dopamine (DA), norepinephrine (Nor), epinephrine (Epi), serotonin (5HT), tyramine (Tyr), and acetylcholine (ACh). WT: *n* = 18 (dopamine, norepinephrine, ACh, tyramine), 17 (epinephrine, serotonin); S123P: n = 12 per agonist. ns = not significant by one-way ANOVA and a Fisher’s LSD. **E.** Sequence alignment of the loop A region. Residue S123 is indicated by an arrow, loop A is highlighted by the orange bar. **F.** AlphaFold2 model of the loop A (orange) region of Dm-DopC1 with S123 relative to loop B (pink), loop C (green) and loop E (blue) at the ligand-binding site.

### 3.5 Cholinergic, dopaminergic and adrenergic compounds antagonise dop-LGICs

To further dissect the pharmacology of the cationic dop-LGICs we tested a range of cholinergic, dopaminergic and adrenergic compounds for their ability to activate or antagonise Dm-DopC1. We found that both Dm-DopC1 dopamine induced currents could be blocked by the nicotinic antagonist tubocurarine, with an IC_50_ of 0.9μM (Figure 6A). In contrast to other nAChRs, the vertebrate a9/10 nAChRs are antagonised by nicotine (Elgoyhen et al., 2001, 1994). In line with this we found that the dopamine currents of Dm-DopC1 could be blocked at high concentrations of nicotine with and IC_50_ of 109μM (Figure 6A). We also tested the muscarinic blocker, atropine, which has been shown to act on nAChRs (Parker et al., 2003; Zwart and Vijverberg, 1997), for its ability to block dopamine currents and found that it behaved similarly to nicotine with relatively poor blocking (IC_50_: 101μM (Figure 6A). This family of dop-LGICs have also been shown to be minimally activated by acetylcholine (Courtney et al., 2025), and due to their close homology with nAChRs and antagonism by nicotine we sought to determine if acetylcholine is able to interact with the dopamine induced current, however we saw no evidence of this (Figure 6A). In addition, we also tested apomorphine, a non-selective vertebrate dopamine receptor agonist and the β-adrenoceptor agonist and antagonist, isoprenaline and propranolol respectively. Apomorphine was able to block the dopamine induced current of Dm-DopC1, with an IC_50_ of 6μM (Figure 6B). Meanwhile propranolol but not isoprenaline was able to block dopamine currents (IC_50_: 100μM) (Figure 6B). We also tested the ability of these compounds to activate the receptors and observed no activation at 100μM (Figure S 3).

**Figure 6.**
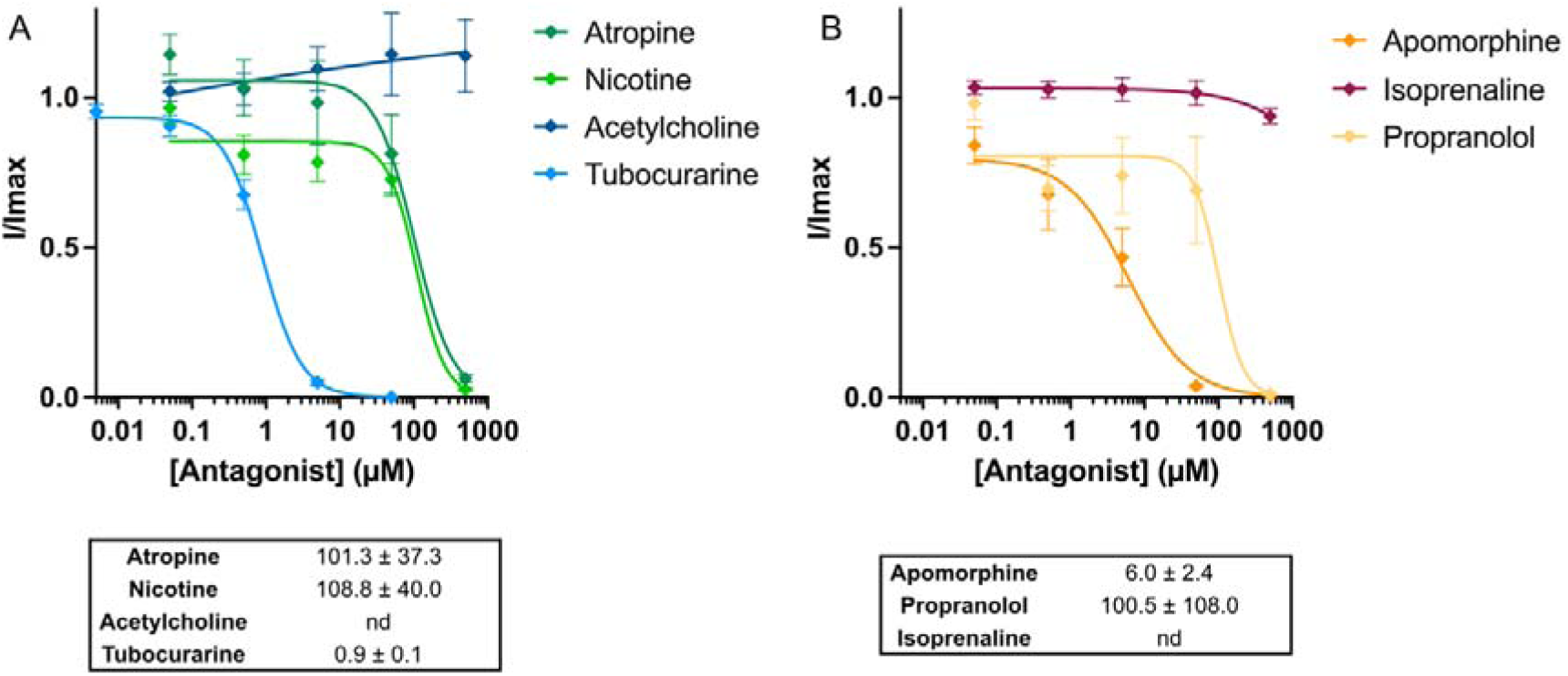
Dm-DopC1 is sensitive to antagonism by cholinergic and dopaminergic compounds. **A-B.** Inhibitory dose response curves for a panel of cholinergic (A) and dopaminergic and adrenergic compounds (B). Error bars represent SEM. *n =* 4 (apomorphine, isoprenaline), 6 (atropine, propranolol), 9 (tubocurarine, acetylcholine), 12 (nicotine). IC50s are shown below each plot and calculated with a 4-parameter curve with constrained bottom = 0, nd = not determined.

### 3.6 *In silico* ligand binding predicts involvement of conserved aromatic cage residues and loop E in ligand recognition

To further predict the binding modality of catecholamines in this novel group of LGICs we employed structural modelling using an AlphaFold3-based co-folding approach (Abramson et al., 2024) in which Dm-DopC1 dimers were co-folded with catecholamines and acetylcholine, 2D interaction predictions were then generated using PoseView (Stierand and Rarey, 2010) (Figure 7). As anticipated in all cases the ligand was predicted to bind within the conserved ligand binding domain composed of the interface between the two adjacent subunits. We also observed that catecholamine binding was predicted to involve many of the highly conserved aromatic cage residues, usually associated with nAChR recognition of acetylcholine. In line with our pharmacological data, dopamine was predicted to have the most direct interactions with amino acids in the binding pocket, including at least one residue from each of the conserved loops (Figure 7A-B). In addition to the aromatic residue interactions dopamine was also predicted to form hydrophobic interactions with L155 of loop E and hydrogen bond with D202 of loop F (Figure 7B). Norepinephrine and epinephrine appeared to have fewer direct contacts, again in line with pharmacological observations in which these ligands have lower receptor sensitivity, however epinephrine maintained an interaction with L155 of loop E (Figure 7C-F). Meanwhile acetylcholine was also predicted to bind in a similar position within the binding pocket however no direct contacts could be predicted (Figure S4).

**Figure 7.**
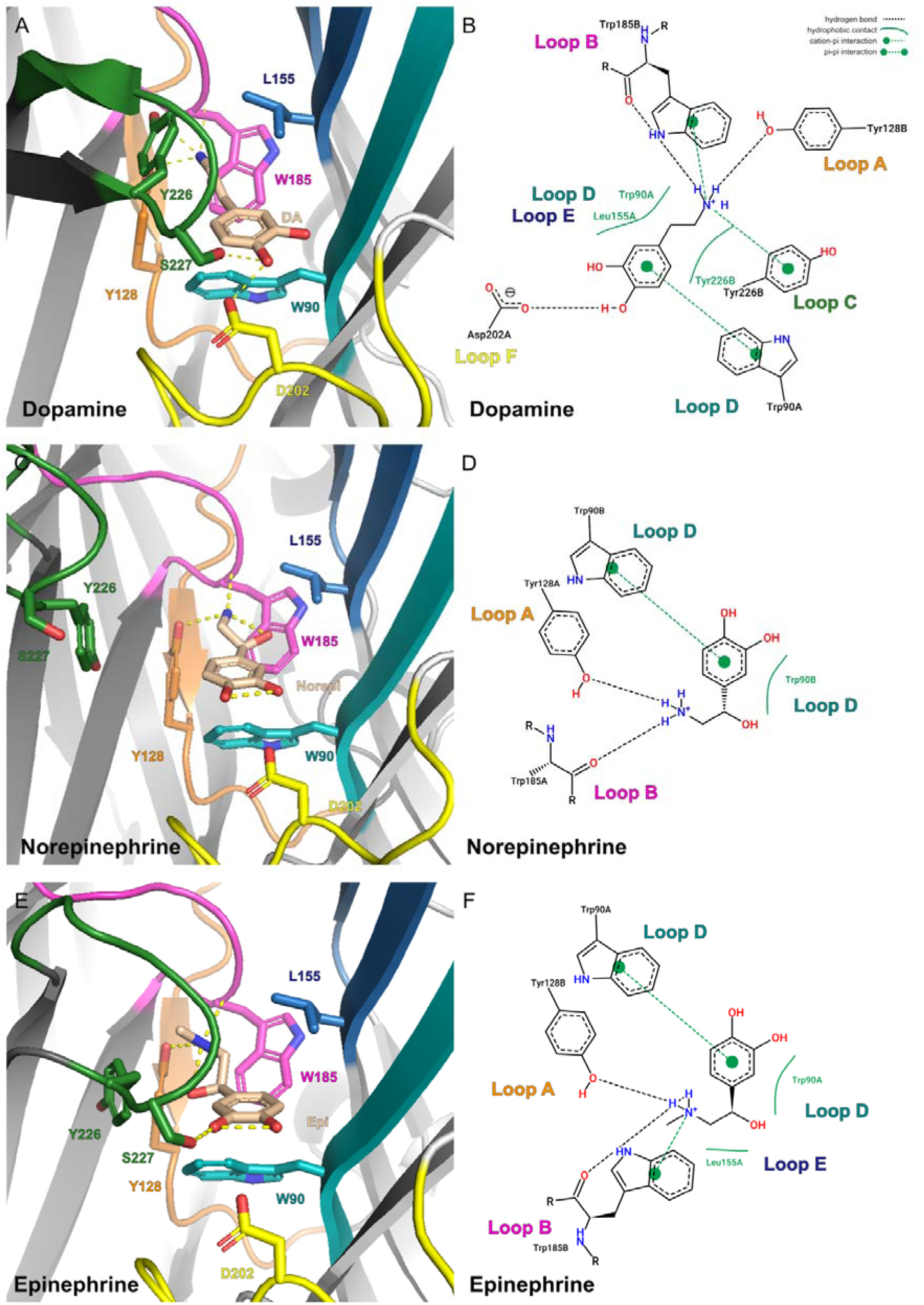
*In silico* binding of catecholamines with Dm-DopC1 predicts involvement of conserved aromatic residues. **A, C & E.** Predicted AlphaFold co-folded models of Dm- DopC1 in complex with dopamine (A), norepinephrine (C) and epinepheline (E). Ligands are shown in beige with polar contacts between the ligand and adjacent residues shown by dashed yellow lines. Colors represent conserved ligand binding loops: A: orange, B: pink, C: green, D: teal, E: blue, F: yellow. Amino acid side chains are shown for residues which appear in 2D poses, as well as S227 which forms a hydrogen bond with dopamine and epinephrine. **B, D & F.** 2D ligand poses for dopamine (B), norepinephrine (D) and epinepheline (F) showing predicted interactions between the ligand and adjacent residues.

## 4. Discussion

Despite high levels of conservation of key ligand binding motifs with vertebrate and invertebrate nAChRs this new class of dopamine-gated cation channels displays striking differences in ligand sensitivity and antagonism by a range of compounds. In this study we highlight some of the key residues that likely contribute to this functional difference (Table 1), with many surrounding previously defined key motifs for cholinergic receptors.

**Table 1.**
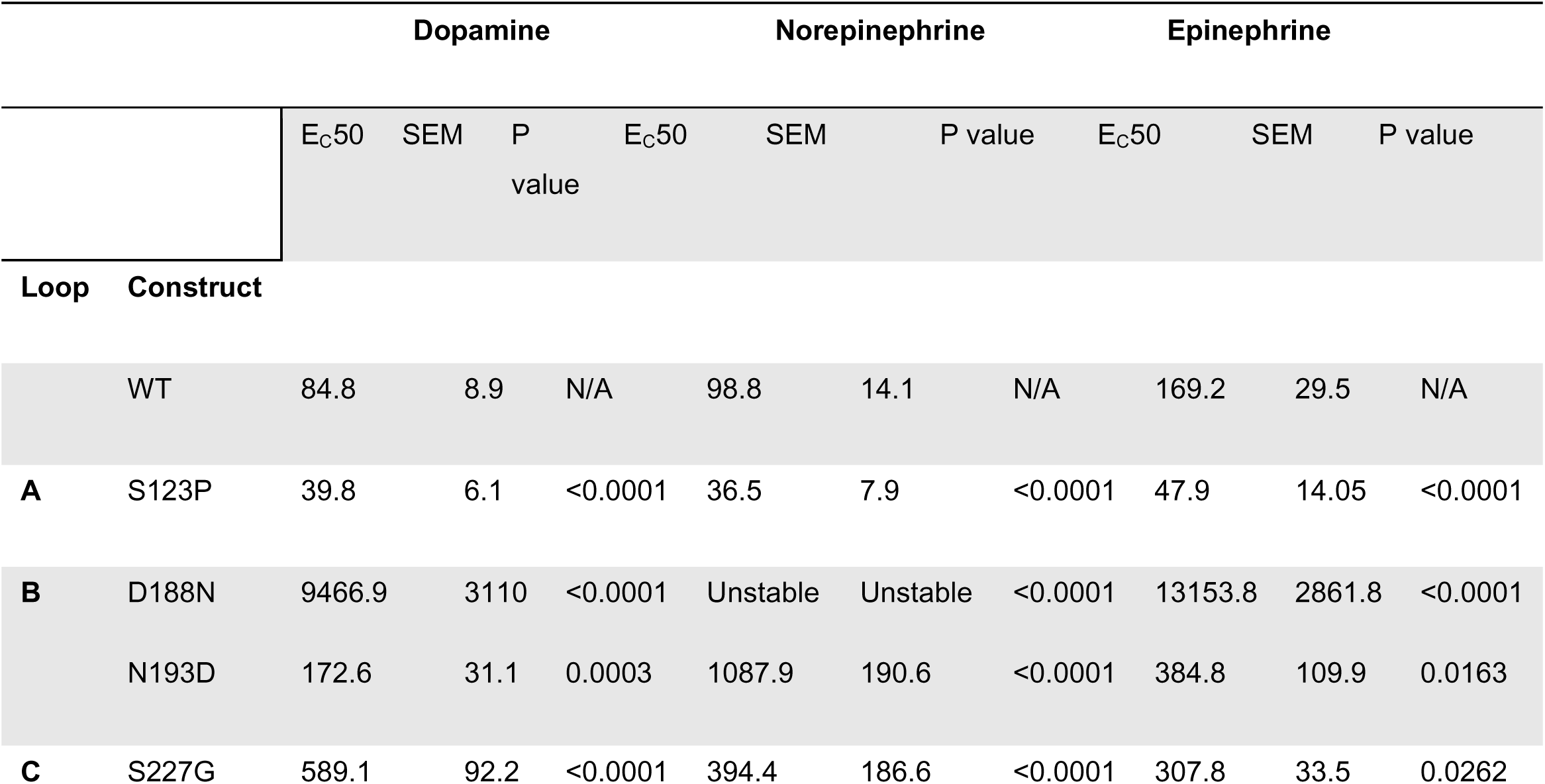
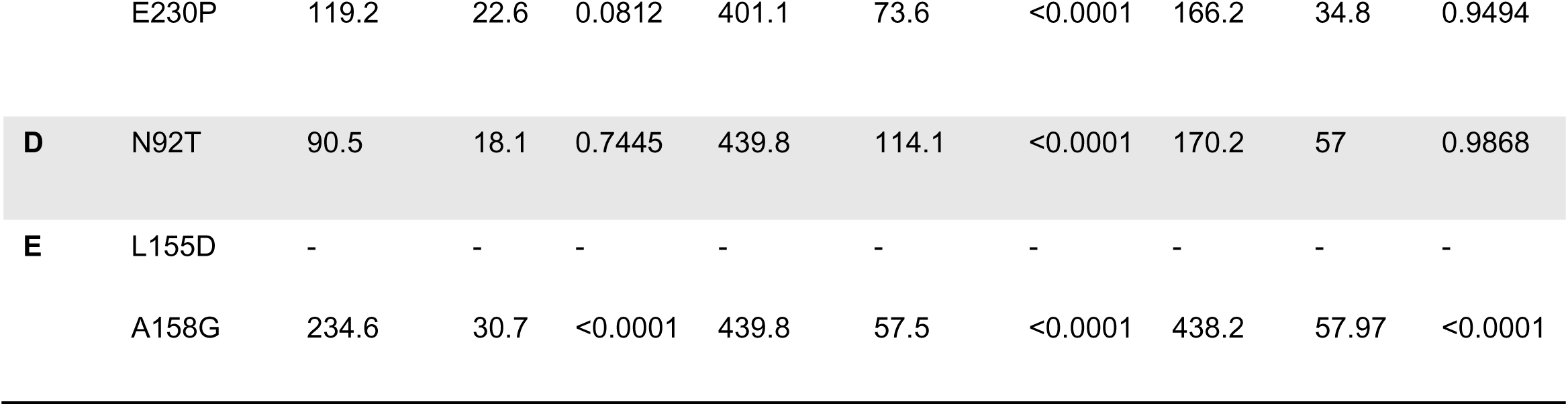
Summary of Ec50 values for all tested mutations. All values are given in μM and calculated using a 3-parameter curve fit with a Hill slope of 1. P values were calculated using the sum of squares F test.

The most striking effect was the almost complete loss of catecholamine gating upon replacement of an aspartate in loop B (D188N) which follows the conserved ‘SWTY’ motif and which is conserved in dop-LGICs. To our knowledge, this site, which in vertebrate α9/10 encodes an asparagine or glycine, has not been previously directly implicated in ligand binding for nAChRs. However, it is unlikely that this residue alone contributes to dopamine selectivity over cholinergic ligands given that aspartate is also found at this position in classic nAChRs such as human α4 and the lack of significant acetylcholine gating over the wild type for this mutation in Dm-DopC1. Our *in silico* ligand binding models also place D188 outside of the core ligand binding residues (Figure S 4), suggesting that it may contributing structurally to the placement of loop B. Interestingly, though immediately following this aspartate is a semi conserved glycine residue which has been shown to distinguish high affinity vertebrate nAChRs (eg. α4 which encodes lysine at this position) from low affinity receptors (eg. α7 which encodes glycine). The increased affinity is thought to be due to stabilization of the open and desensitized states by virtue of a hydrogen bond between this position in loop B and a proline in loop C that forms in absence of glycine (Grutter et al., 2003; Kouvatsos et al., 2016; Puskar et al., 2012). Interestingly glycine is present at this position in all but one dop-LGIC (which instead encodes an alanine) further supporting the hypothesis that this hydrogen bond is key to high affinity acetylcholine receptors. Decoupling of loops B and C may be important for gating by aminergic ligands given that monoamine gated receptors from other subfamilies such as the vertebrate 5HT_3_R and invertebrate chloride permeable dopamine and histamine receptors lack the hydrogen forming proline within loop C. In addition, loop B also contains one of the conserved tryptophans associated with aromatic cage formation. Preceding this tryptophan by 2 amino acids is another glycine residue conserved in nAChRs α subunits but encoding an arginine in β subunits, which causes movements in the tryptophan preventing it from forming a functional binding site (Morales-Perez et al., 2016), in dop-LGICs this site encodes a alanine or serine. Mutations in loop B have also been shown to be more broadly required for monoamine binding in 5HT_3_Rs for example (Thompson et al., 2008). It would be interesting to see if the D188N reverse mutation in vertebrate receptors alters acetylcholine gating, although given the divergence of sequences at this position across nAChRs in both vertebrates and invertebrates this residue may not be contributing in a significant way to acetylcholine recognition.

We also confirmed the importance, yet surprising flexibility, of a highly conserved proline residue within the ‘WXPD’ motif of loop A, which has been hypothesized to be critical for maintaining structural integrity and receptor assembly in pLGIC receptors from bacteria through to humans (Braun et al., 2016). We observed that Dm-DopC1, which exhibits a higher EC_50_ for dopamine than other dop-LGICs (Courtney et al., 2025), diverged from this motif to encode serine at this position (WKSD), which when substituted for proline, led to significant increases in catecholamine sensitivity. The mirrors previous work in glycine receptors where the same proline was replaced by an alanine leading to a decrease in ligand sensitivity (Hughes et al., 2020; Lummis and Dougherty, 2022). Not only does our finding further highlight the importance of a proline at this position, but it also shows that there is flexibility in the motif, as the natural variant still displays micromolar ligand binding. As this serine is not conserved within the rest of the dop-LGIC group, it is unlikely to be directly participating in catecholamine recognition. Rather this suggests, as has been previously hypothesized, that the conserved proline of the ‘WXPD’ motif contributes indirectly by stabilizing loop A conformation or facilitating ligand alignment within the binding pocket.

An intriguing feature of the invertebrate cationic dop-LGICs is the presence of key motifs usually attributed to acetylcholine and nicotine binding in nAChRs. This is highlighted by our *in silico* binding experiments which suggest that many of the key residues involved in aromatic cage formation, usually associated with acetylcholine binding participate in the binding of dopamine and the other catecholamines. In particular this can be seen in loop C, which in dop-LGICs also encodes the conserved ‘YXCC’ motif (Karlin, 2002; Tomizawa et al., 2008) as well as a pair of tyrosines implicated in aromatic cage formation (Braun et al., 2016). However, the at ‘X’ position of the ‘YXCC’ motif the dop-LGICs encode a conserved serine, which is not present in nAChRs. By exchanging this serine for glycine (present in α9/10) in Dm-DopC1, we found that this serine is indeed important for catecholamine gating but did not confer acetylcholine gating despite restoring the motif often found in nAChRs. Interestingly our binding pose predictions did not identify direct interactions of the ligands with this serine, however for both dopamine and epinephrine a hydrogen bond between serine and the ligand was predicted. Separately, we also found that the residue immediately downstream of the ‘YXCC’ motif was important for determining differences in ligand sensitivity between *Daphnia* dop-LGICs. The functional impact of mutations at these positions supports the established role of loop C in maintaining the geometry of the ligand binding pocket, particularly in orienting conserved aromatic residues that contribute to catecholamine recognition. It also echoes our findings within loop B, which suggests that a series of changes in the binding regions are required to completely alter receptor selectivity from cholinergic to dopaminergic.

We also explored the role of loops D, E and F that form the complimentary face of the ligand binding pocket and have been shown to contribute to stabilizing the extracellular domain. A key feature of Loop D is the ‘WXD’ motif that is conserved across nAChRs, dop-LGICs, 5HT_3_Rs and even some chloride channels (Braun et al., 2016; Corringer et al., 1995; Spier and Lummis, 2000). Loop D is remarkably well conserved between dop-LGICs and nAChRs, suggesting that its main function is to stabilize the ligand binding regions. However, we did note that both Dm-DopC2 and Am-DopC1, which are more selective for dopamine than the other dop-LGICs encoded a threonine 2 amino acids before the ‘WXD’ motif, rather than an asparagine. When we substituted this asparagine for threonine in Dm-DopC1 we noted a specific alteration in ligand sensitivity with only the norepinephrine EC_50_ being adversely affected while dopamine and epinephrine responses remained like wild type, making the receptor overall more selective for dopamine. Interestingly, vertebrate nAChRs typically encode a lysine or arginine at this position which is also conserved across other pLGICs including glycine and GABA receptors, where disruption of this residue leads to almost complete loss of ligand activation (Goldschen-Ohm et al., 2011; Grudzinska et al., 2005). The cationic dop-LGICs, in line with hypotheses drawn up by Lynagh and Pless in 2014, also lack positively charged residues in this position instead favoring polar residues also found in other amine gated receptors including HisCl and the anionic dop-LGICs from nematodes, which may assist in binding the hydroxyl termini or these ligands (Lynagh and Pless, 2014). In addition, our *in silico* model also predicted direct hydrophobic interactions between L155 of loop E and dopamine and epinephrine. Hydrophobic residues at this position are conserved across dop-LGICs in contrast to vertebrate α9/10 nAChRs which encode a charged aspartate at this position. Interestingly when we exchanged leucine for aspartate at this position, we could not generate significant dose dependent ligand-induced currents for any of the catecholamines nor acetylcholine. This is mirrored by a previous study of Am-DopC1 which identified the adjacent serine residue, which is conserved across dop-LGICs, as critical for dopamine binding (Mitchell et al., 2025). Taken together these data suggest this region of loop B is important for ligand recognition in dop-LGICs. Collectively these mutations have identified residues critical for catecholamine recognition across various conserved ligand binding domains, revealing that like in other pLGICs, both faces of the ligand binding domain are critical not only for ligand gating but also relative ligand selectivity. However, this study also demonstrates that ligand switching from acetylcholine to (or from) dopamine requires a series of alterations across evolutionary time and across multiple regions of the binding domain.

Finally, we also examined the pharmacology of dop-LGICs by testing a range of cholinergic, dopaminergic and adrenergic compounds for their ability to interact with the dopamine induced currents from Dm-DopC1. The competitive nAChR antagonist, tubocurarine (Rahman et al., 2022), efficiently blocked currents in the low μM range, broadly in line with classic nAChRs (Zwart and Vijverberg, 1998). In contrast, other cholinergic compounds such as nicotine and atropine (a muscarinic antagonist) had effects only at high concentrations. This is in line with evidence that tubocurarine can act on a broader range of pLGICs including vertebrate 5HT_3_Rs (Peters et al., 1990; Yan et al., 1998) and invertebrate chloride gated cholinergic receptors (Hardege et al., 2023). In addition, dop-LGICs could be blocked by apomorphine (non-selective vertebrate dopamine agonist) and propranolol (β-adrenoceptor antagonist) in the μM range suggesting that dop-LGICs have a distinct pharmacology from their closely related nAChRs, with the potential of novel chemistries to be used to target this subfamily.

## 5. Conclusion

In this study we detail the mechanistic map of ligand selectivity of the invertebrate dopamine-gated cation channel (dop-LGIC) family, extending the known functional repertoire of these receptors. Our primary aim was to identify how variations in conserved ligand binding domains enable dop-LGICs to respond selectively to catecholamines rather than cholinergic ligands, despite their close sequence homology with vertebrate nAChRs. In line with this we found that while core structural features of the pLGIC superfamily are conserved, functional divergence is driven by key residue variations surrounding these canonical motifs in the extracellular loops. We identified residues critical for catecholamine recognition across loops typically associated with both the principal (A-C) and complementary faces (D-F) of the ligand binding domain. Pharmacological analysis of dop-LGICs also confirmed a distinct profile from their nAChR relatives, with dopamine-induced currents efficiently blocked in the micromolar range by cholinergic, dopaminergic and adrenergic compounds.

Our results provide insight into the functional plasticity within invertebrate neurotransmission systems and reveal how subtle differences in loop composition enable the switch from cholinergic to aminergic ligand recognition. The fact that cationic dop-LGICs are conserved across multiple invertebrate species, confirms the widespread nature of this signaling mechanism. Furthermore, the demonstrated requirement for a series of evolutionary changes to achieve ligand switching raises fundamental questions about the origin of ionotropic neurotransmission, specifically whether dopamine or acetylcholine signaling emerged first. Future research should focus on comparative structural analysis of dop-LGICs across basal phyla, such as Platyhelminths or Nematodes (where unique anionic dop-LGICs are already found), to trace the evolutionary trajectory and structural requirements that drove the transition between cholinergic and dopaminergic ionotropic signaling.

## Acknowledgements

The authors kindly thank Amy Courtney & William Schafer for providing the plasmid containing wild type Dm-DopC1. The authors also gratefully acknowledge Howard Baylis and all present and past members of the Hardege lab for helpful discussions.

## Author Contributions

**T. M. M:** Conceptualization, Methodology, Validation, Formal analysis, Investigation, Data Curation, Writing - Original Draft, Visualization.
**T. E. R:** Methodology, Validation, Formal analysis, Investigation, Data Curation, Writing - Review & Editing.
**Y. Z:** Formal analysis, Investigation, Data Curation.
**T. R:** Supervision, Project administration.
**I. H:** Conceptualization, Formal analysis, Resources, Data Curation, Writing - Original Draft, Visualization, Supervision, Project administration, Funding acquisition.

## Supplementary data

**Table S1.**
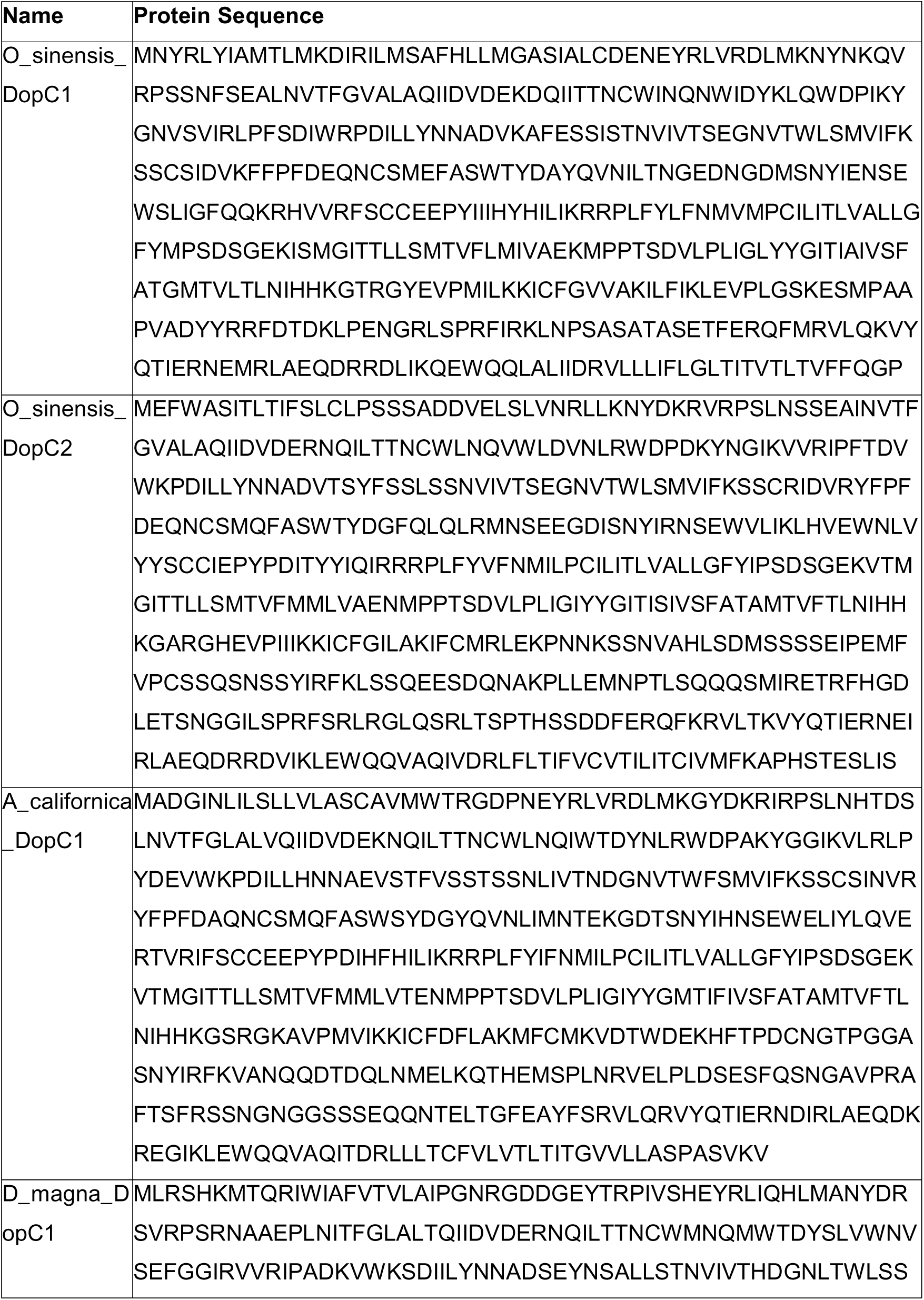
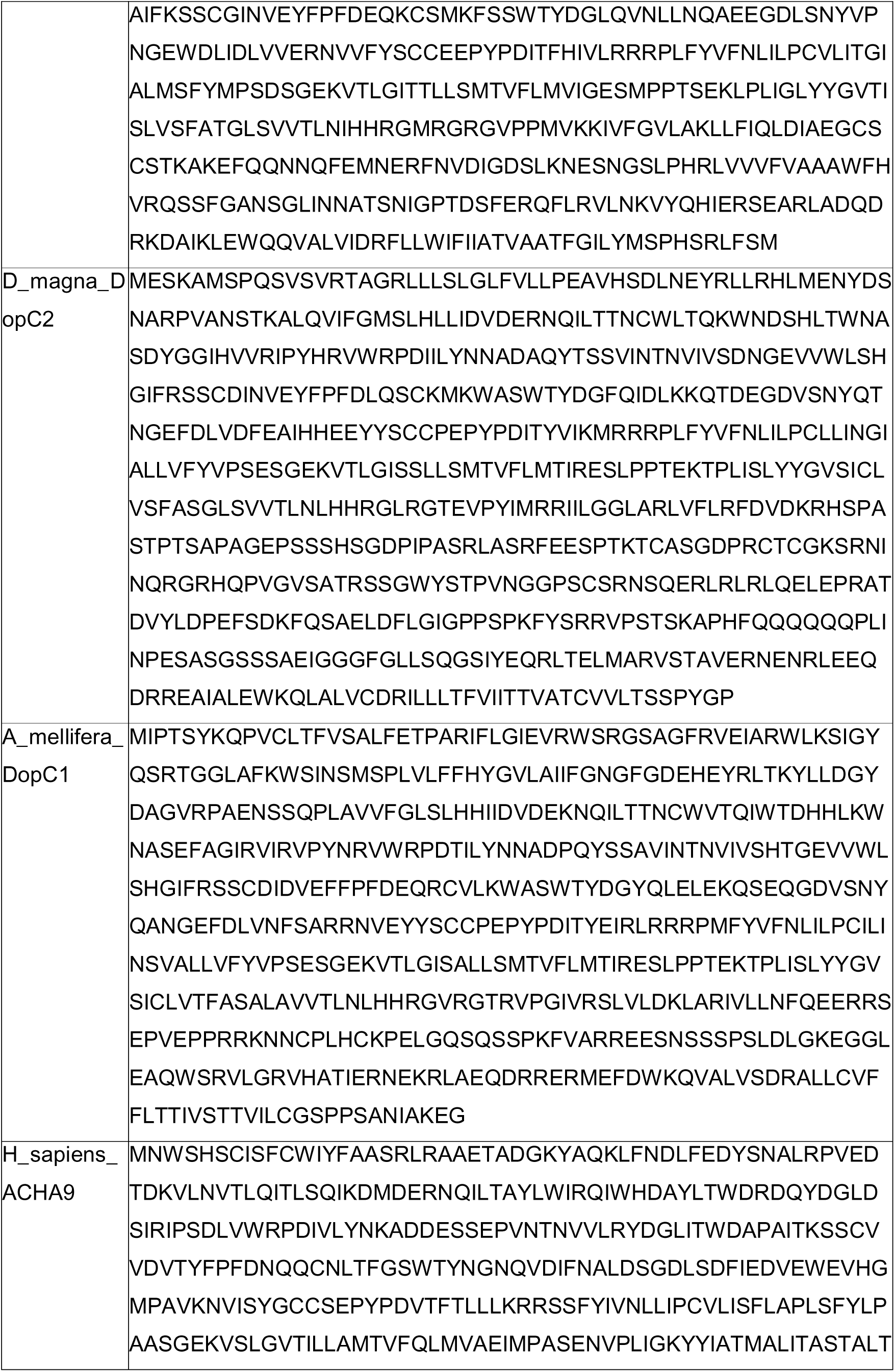
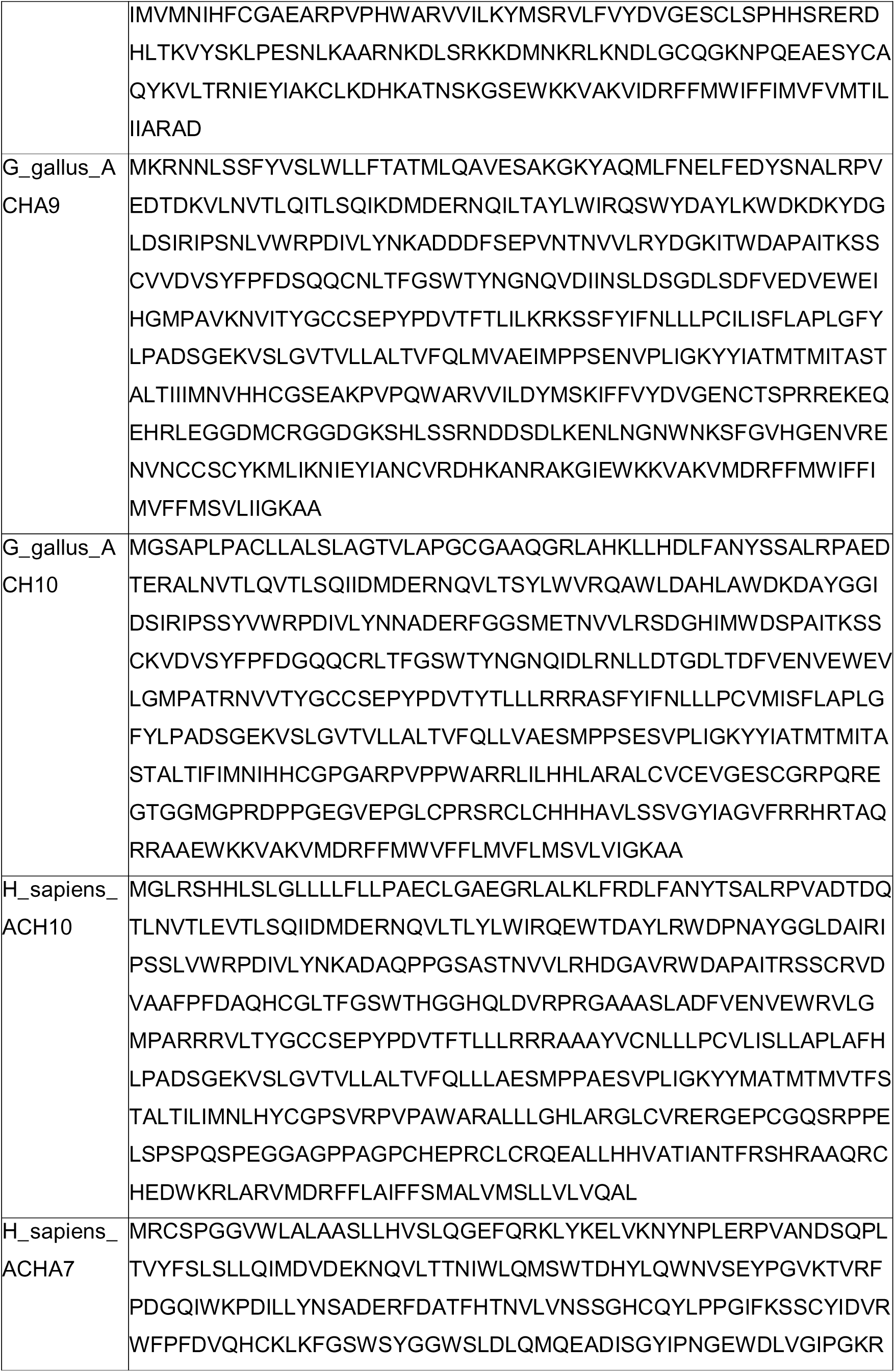
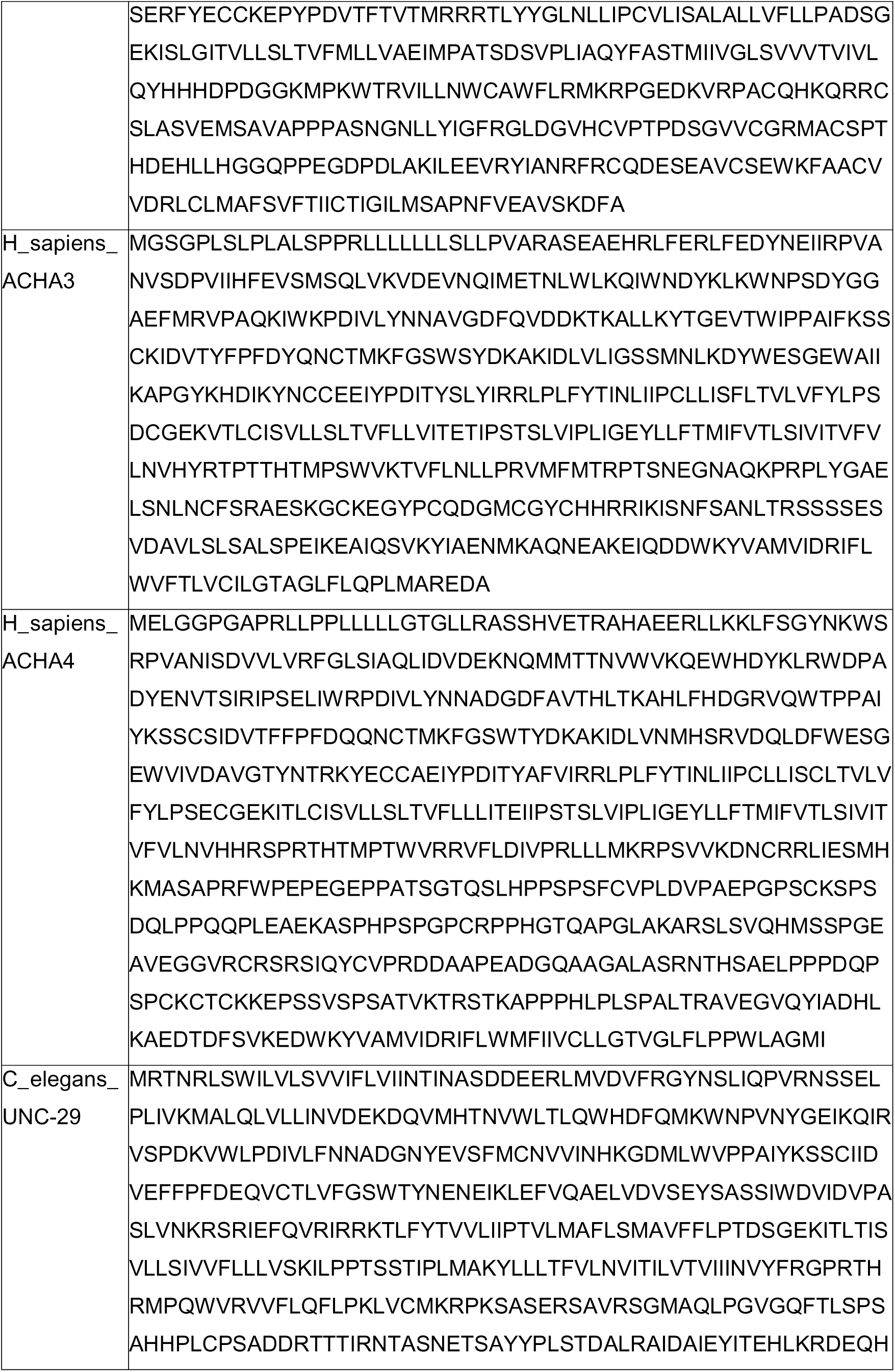
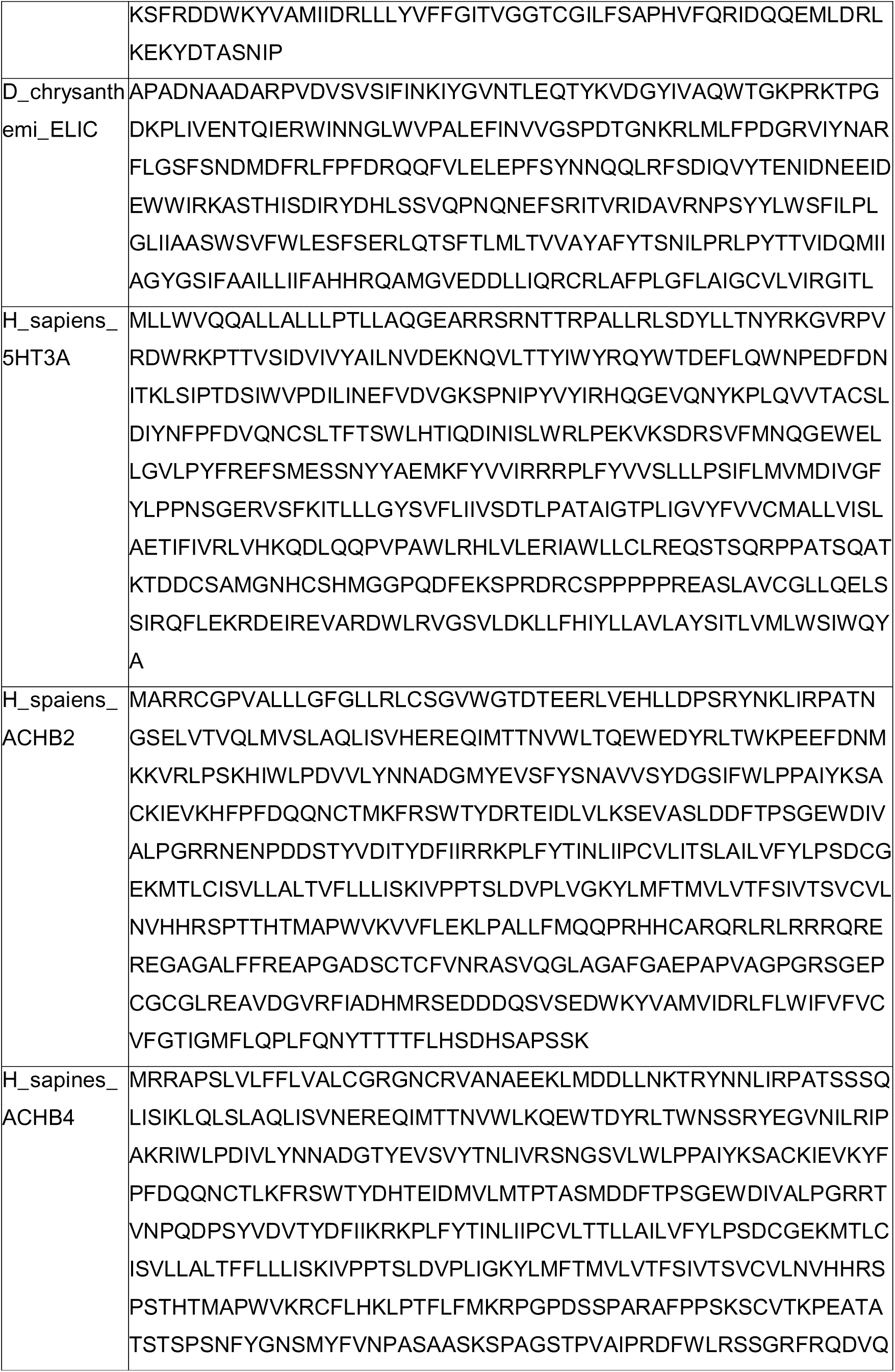
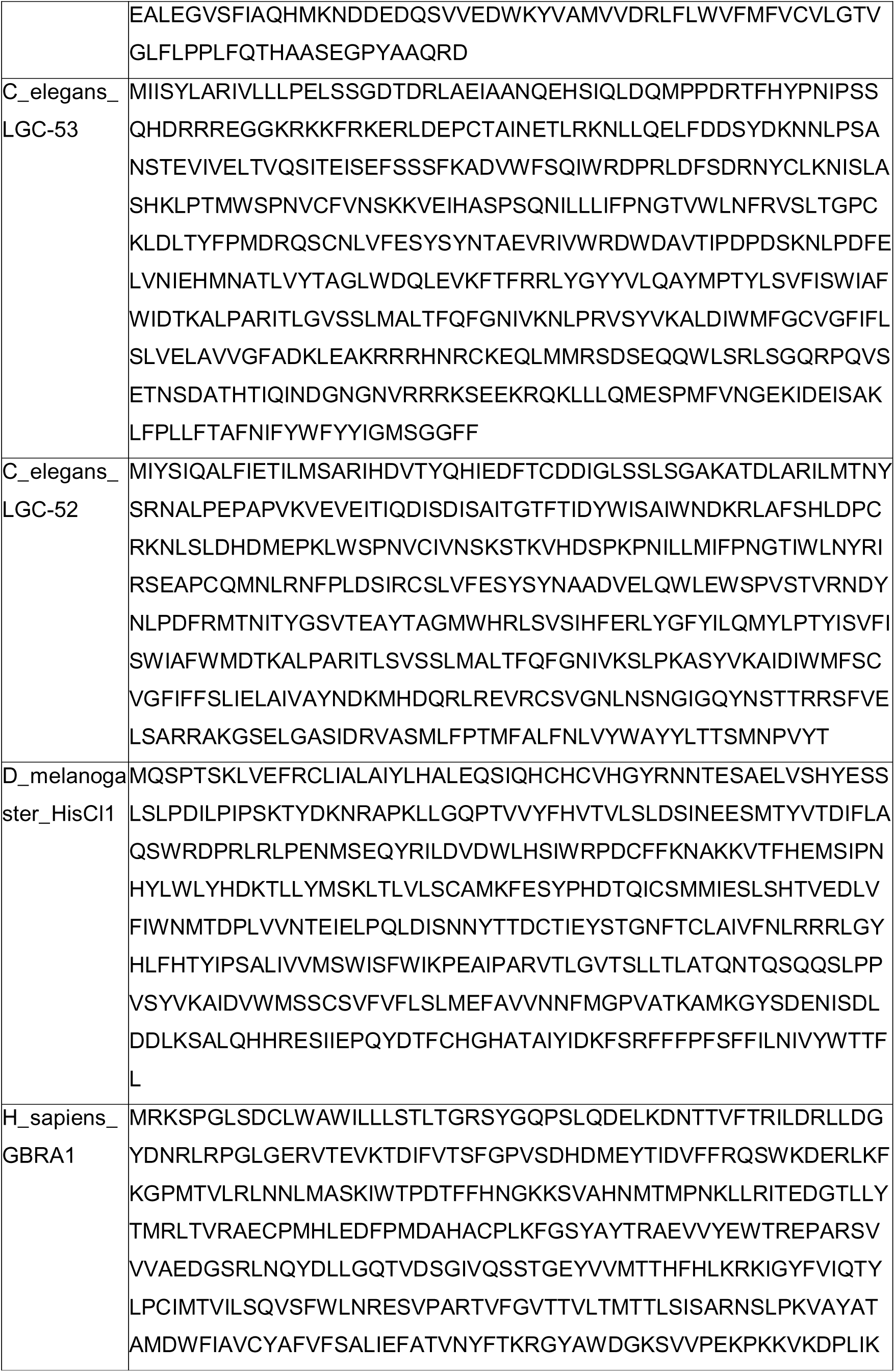
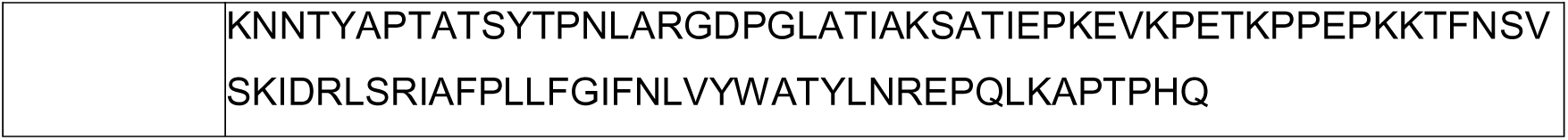
Protein sequences used for alignments.

**Figure S1.**
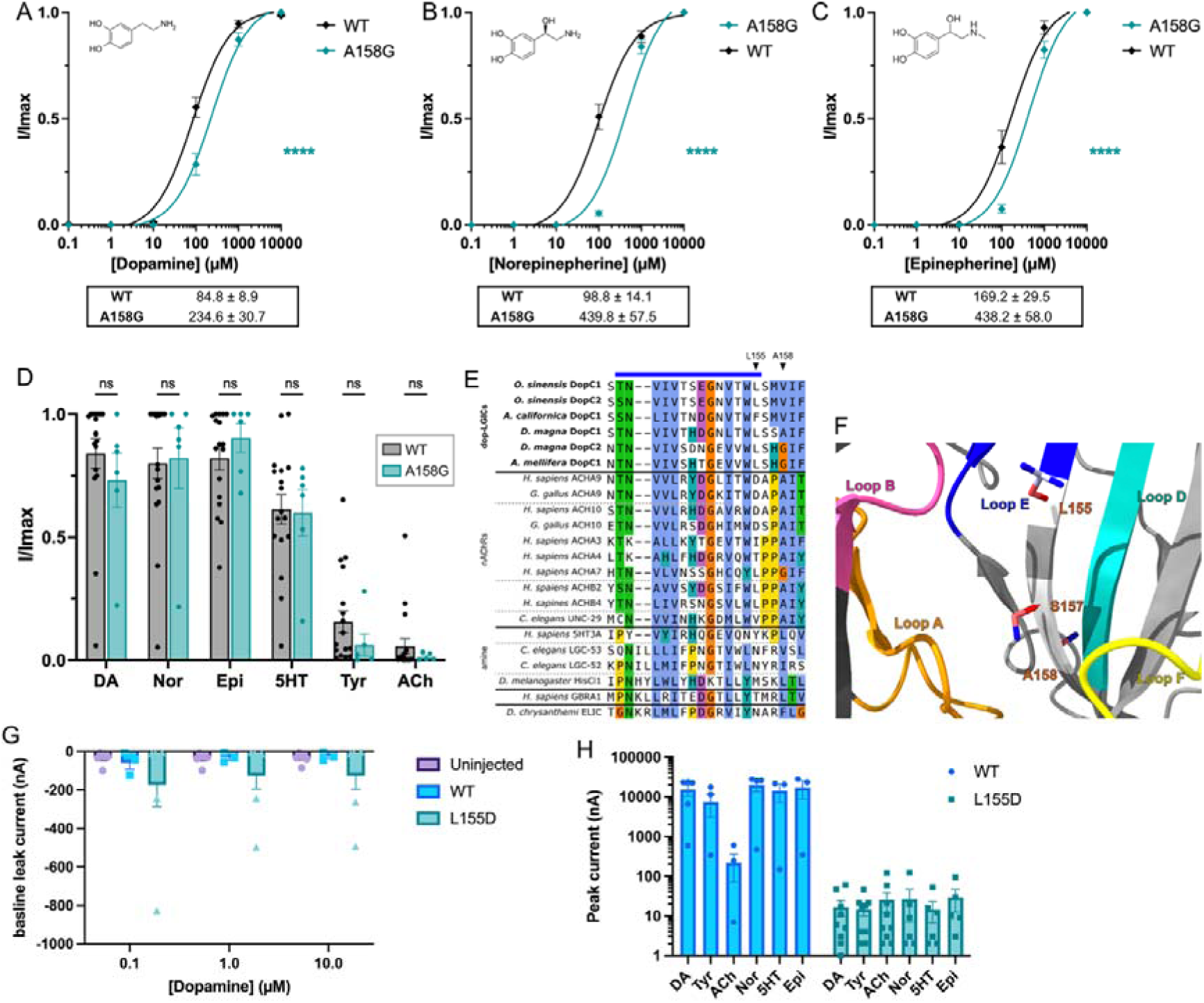
The A158G mutation in Loop E of a *D. magna* dop-LGIC decreases catecholamine sensitivity. **A-C.** Dose response curves for dopamine (A), norepinephrine (B) and epinephrine (C) in wild-type and A158G Dm-DopC1 expressing *Xenopus* oocytes. Current normalized by I/Imax. EC_50_ values (± SEM) shown below each plot. **** = P<0.0001, ns = not significant calculated by sum of squares F test. WT: *n* = 24 (dopamine), 12 (norepinephrine), 11 (epinephrine); A158G: *n* = 10 (dopamine), 11 (norepinephrine), 10 (epinephrine). D. Relative maximum I/Imax currents of *Xenopus* oocytes expressing wild-type and A158G Dm-DopC1, perfused for 10 s with 1 mM each: dopamine (DA), norepinephrine (Nor), epinephrine (Epi), serotonin (5HT), tyramine (Tyr), and acetylcholine (ACh). WT: *n* = 18 (dopamine, norepinephrine, ACh, tyramine), 17 (epinephrine, serotonin); A158G: *n* = 6 per agonist. ns = not significant by one-way ANOVA and a Fisher’s LSD. E. Sequence alignment of the loop E region. Residues A158 and L155 are indicated by arrows, loop E is highlighted by the blue bar. F. AlphaFold2 model of the loop E (blue) region of Dm-DopC1 with A158 and L155 relative to loop B (pink), loop C (green), loop F (yellow), loop A (orange) and loop D (turquoise) at the ligand-binding site. G. Baseline currents in uninjected oocytes or oocytes expressing WT or L155D Dm-DopC1. Baseline currents measured 5s before agonist application. Error bars represent SEM, *n =* 3 (WT), 6 (uninjected), 7 (L155D). H. Peak currents in response to 10s agonist application of *Xenopus* oocytes expressing wild-type and L155D Dm-DopC1 with 1mM each: dopamine (DA), norepinephrine (Nor), epinephrine (Epi), serotonin (5HT), tyramine (Tyr), and acetylcholine (ACh). Error bars represent SEM. WT: *n* = 5 (dopamine), 4 (norepinephrine, ACh, tyramine), 3 (epinephrine, serotonin); L155D: *n* = 9 (dopamine, tyramine, ACh), 6 (norepinephrine, serotonin), 5 (epinephrine). Note scale bar on log_10_ scale.

**Figure S2.**
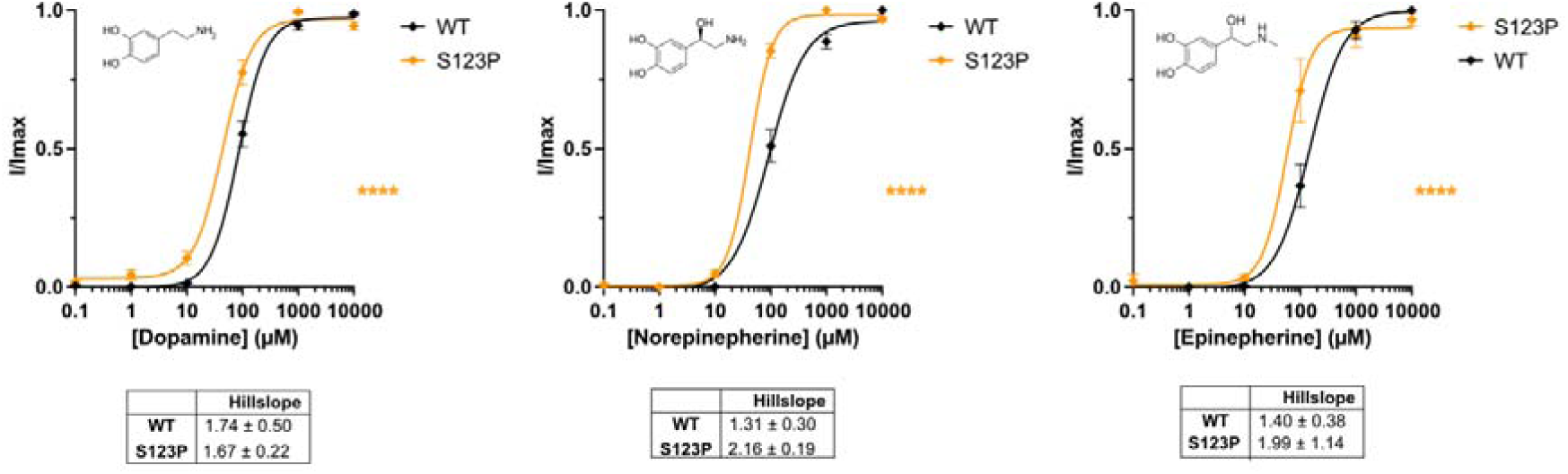
S123P exhibits a steeper Hill slope than wild type for norepinephrine. **A-C.** Dose response curves for dopamine (A), norepinephrine (B) and epinephrine (C) in wild-type and S123P Dm-DopC1 expressing *Xenopus* oocytes. Current normalized by I/Imax. Hill slope values are shown underneath each plot with SEM, calculated with a 4-parameter curve. WT: *n* = 24 (dopamine), 12 (norepinephrine), 11 (epinephrine); S123P: n = 12, (dopamine), 4 (norepinephrine) and 8 (epinephrine).

**Figure S3.**
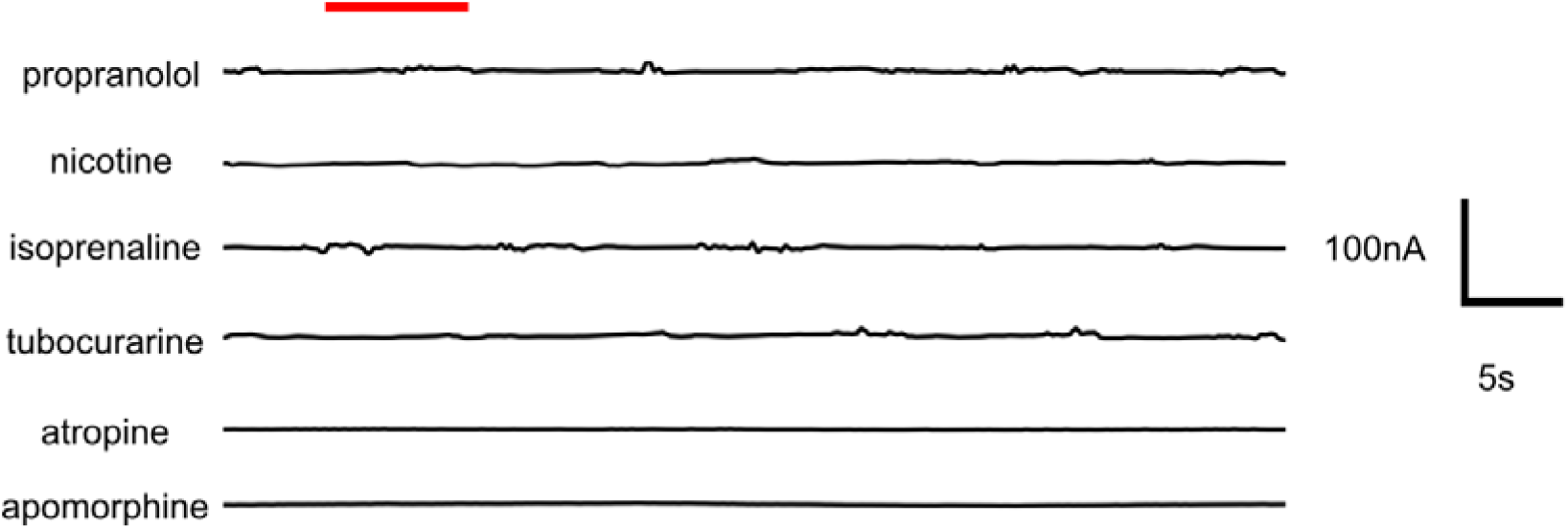
Continuous traces of Dm-DopC1 oocytes exposed to a panel of compounds. Red bar indicates application of 100μM compound.

**Figure S4.**
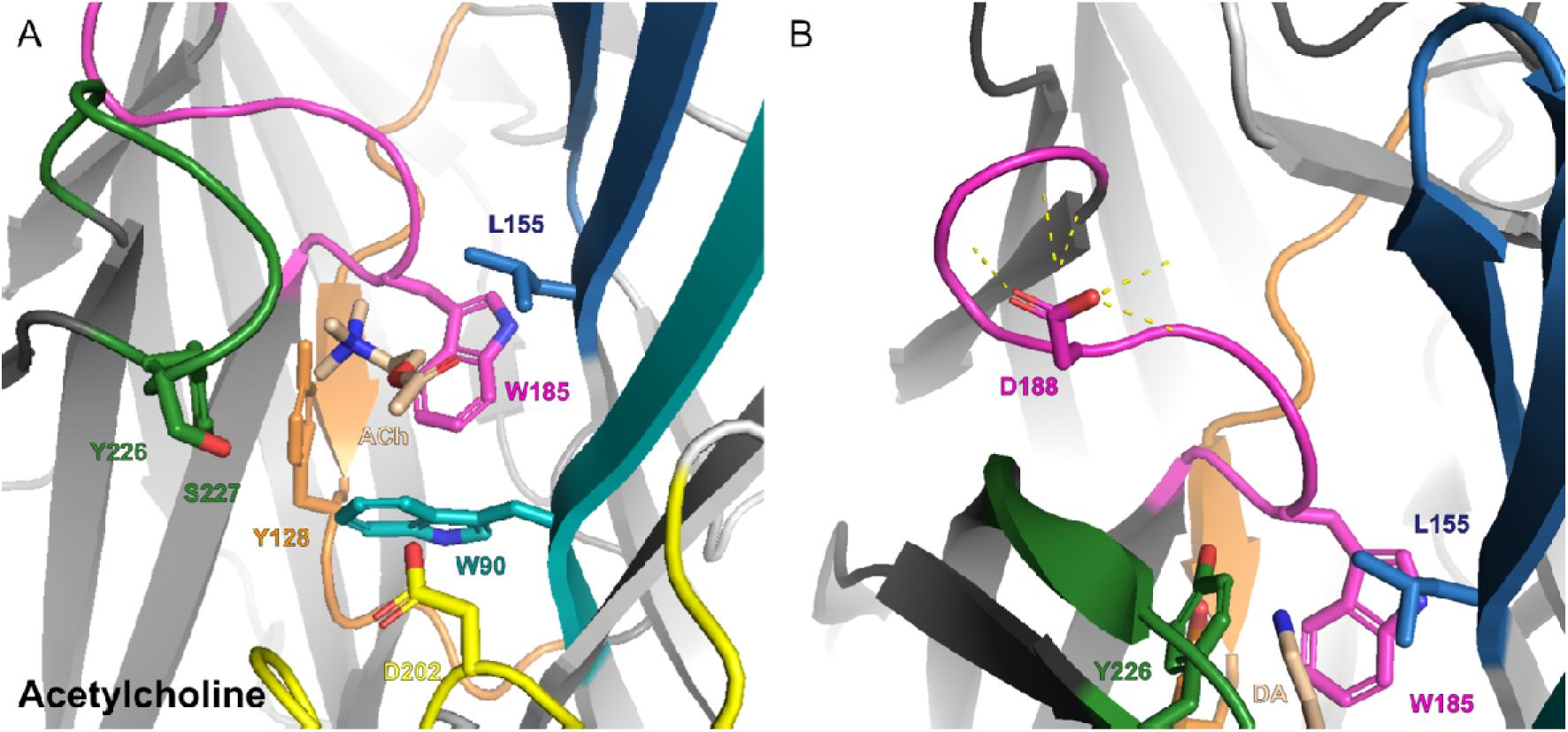
*In silico* binding models of Dm-DopC1 with acetylcholine (A) and dopamine (B). **A & B**. Predicted AlphaFold3 co-fold models of Dm-DopC1 in complex with acetylcholine (A) and dopamine (B). Ligands are shown in beige with polar contacts between the ligand and adjacent residues shown by dashed yellow lines. Colors represent conserved ligand binding loops: A: orange, B: pink, C: green, D: teal, E: blue, F: yellow. Amino acid side chains are shown for residues which appear in 2D poses for dopamine, as well as D188 and S227 which forms a hydrogen bond with dopamine.

